# Genomic data reveals habitat partitioning in massive *Porites* on Guam, Micronesia

**DOI:** 10.1101/2024.07.16.603743

**Authors:** Karim D. Primov, David R. Burdick, Sarah Lemer, Zac H. Forsman, David J. Combosch

**Affiliations:** University of Guam Marine Laboratory, UOG Station, Mangilao, GU, USA; Department of Integrative Biology, University of Texas at Austin, Austin, TX, USA; King Abdullah University of Science and Technology, Thuwal, 23955, Saudi Arabia

**Keywords:** Massive *Porites*, coral, phylogenomics, population genomics, marginal river deltas, Guam, ddRAD

## Abstract

Corals in marginal reef habitats generally exhibit less bleaching and associated mortality compared to nearby corals in more pristine reef environments. It is unclear, however, if these differences are due to environmental differences, including turbidity, or genomic differences between the coral hosts in these different environments. One particularly interesting case is in the coral genus *Porites*, which contains numerous morphologically similar massive *Porites* species inhabiting a wide range of reef habitats, from turbid river deltas and stagnant back reefs to high-energy fore reefs. Here, we generate ddRAD data for 172 *Porites* corals from river delta and adjacent fore reef populations on Guam to assess the extent of genetic differentiation among massive *Porites* corals in these two contrasting environments and throughout the island. Phylogenetic and population genomic analyses identify seven different clades of massive *Porites*, with the two largest clades predominantly inhabiting either river deltas and fore reefs, respectively. No population structure was detected in the two largest clades, and *Cladocopium* was the dominant symbiont genus in all clades and environments. The perceived bleaching resilience of corals in marginal reef environments may therefore be attributed to interspecific differences between morphologically similar species, in addition to potentially mediating environmental differences. Marginal reef environments may therefore not provide a suitable refuge for many reef corals in a heating world, but instead host additional cryptic coral diversity.

## Introduction

Coral taxa living in sub-optimal or marginal environmental conditions may prove resistant to both global and local stressors. Turbid river delta reefs, for example, are generally characterized as sub-optimal environments for corals due to higher nutrient content, sedimentation rates, and turbidity compared to most other reef types, especially oceanic fore reefs [1,2]. These reefs, however, have been identified as potential refuges for threatened marine taxa [2,3] because marginal reef populations can withstand their native habitat’s suboptimal conditions [3–7]. Indeed, marginal reef corals have exhibited less bleaching and associated mortality compared to nearby shallow-water reefs at multiple locations throughout the Indo-Pacific [8–10].

Such resilience to environmental stressors, including differential bleaching response among corals, may either be environmentally mediated, driven by interspecific or intraspecific genetic differences [2,11], or the result of genotype-environment interactions. Two of many environmental factors influencing coral survival and resilience during bleaching events in river deltas are turbidity and heterotrophy, factors which dominant marginal reef coral taxa may have likely adapted to [2]. Furthermore, coral host population structure and differences in protein profiles between nearshore and offshore corals have been identified for both Caribbean and Indo-Pacific *Porites* [12–14]. It is still unclear, however, whether differential bleaching response between marginal and oceanic reef corals can manifest solely from population-level genomic differences or can also be attributed to species-level genomic differences or plasticity.

Resolving the underlying factors manifesting in differential bleaching response is confounded by morphological similarity and phenotypic plasticity of closely related yet genetically distinct species. Both of these factors are widespread among marine taxa, especially in reef-building corals [15], and has led to difficulties in properly identifying coral taxa, leading to large knowledge gaps with respect to both geographic and ecological distributions of specific coral species [16–21]. This is especially the case for corals in the genus *Porites* [17,22–24]. *Porites* is one of the most prominent reef-building coral genera throughout the Indo-Pacific. Within the genus, massive *Porites*, a dominant, cosmopolitan ecoform of corals consisting of an unresolved number of species, have been found to be resilient to a variety of environmental stressors, including low pH, thermal stress, and turbidity [16,17,22–31].

The Pacific island of Guam is host to a variety of massive *Porites* species, such as *P. australiensis, P. densa*, *P. lobata*, *P. lutea*, *P. murrayensis*, *P. stephensoni*, and *P. solida* [32–34]. Massive *Porites* on Guam can be found in a variety of different habitats, ranging from oceanic fore reefs to marginal turbid river deltas, where these contrasting habitats can be found less than 0.5 km from one another in Southern Guam. River deltas on Guam are geographically confined to the southern region of the island, which is composed of volcanic terrane, while northern Guam is composed mainly of porous limestone (Figure 1) [35,36]. Massive *Porites* in these two contrasting habitats in Southern Guam provide the perfect study system to identify whether differential bleaching response is driven by either environmental differences or interspecific or intraspecific genomic differences.

**Figure 1:**
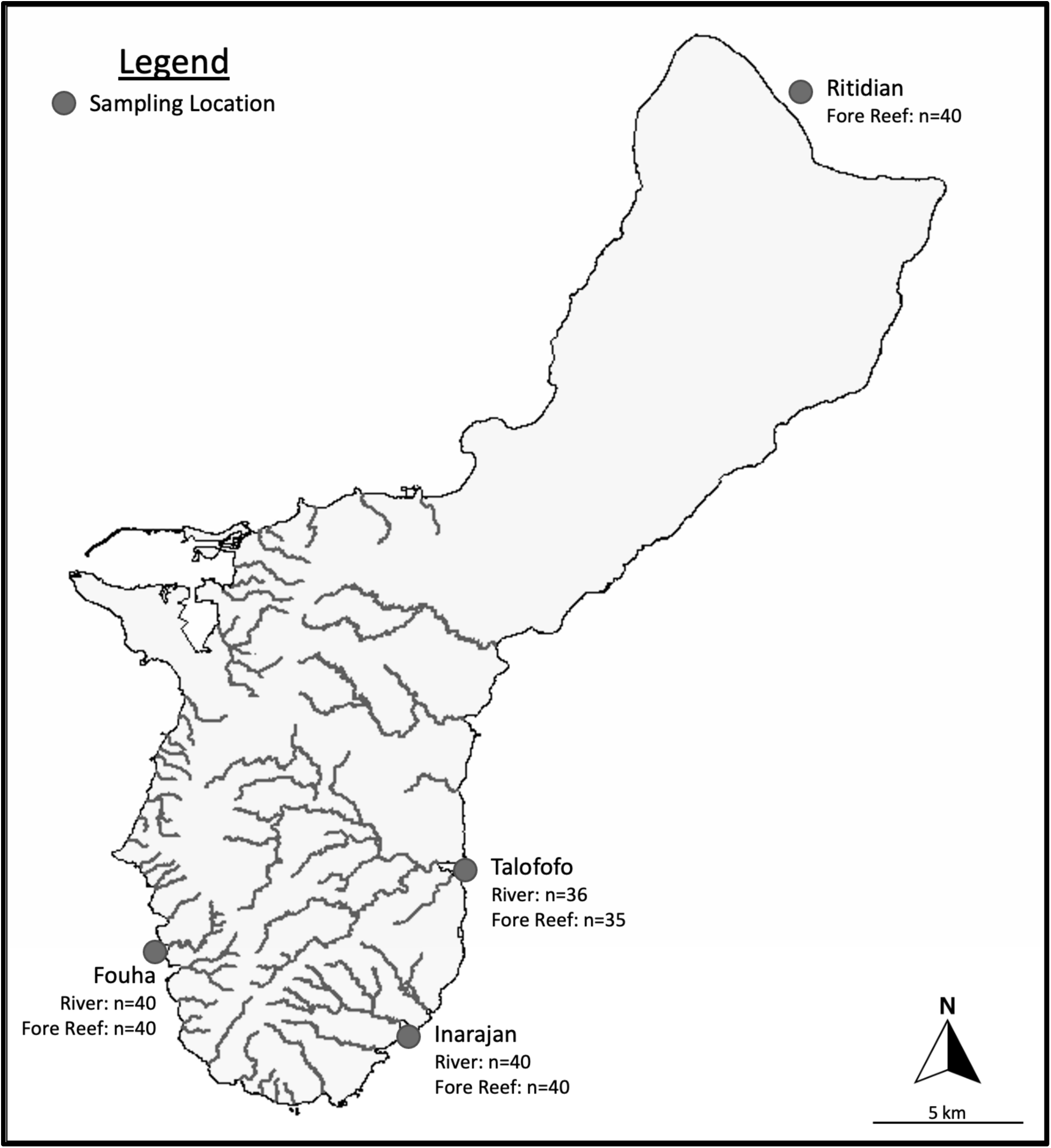
Geographic extent of sampling of massive Porites on Guam (Credit: Pacific Island Data Portal). Lines correspond to rivers and tributaries, which are found exclusively in Southern Guam. Overall, 271 massive Porites samples were collected for this study but only 172 were genotyped and analyzed morphologically.

In this study, we employ double-digest RADSeq (ddRAD) to determine whether genetic differentiation exists between river delta and adjacent massive *Porites* populations in Southern Guam, and whether existing genetic differences between corals in these contrasting habitats are interspecific or intraspecific. We further assess basic population genomic characteristics among major massive *Porites* lineages, both between these contrasting habitats and around Guam.

## Materials and Methods

### Site selection and sample collection

A total of seven populations of massive *Porites* corals were sampled around Guam (Table S1). Three river delta populations were sampled at Inarajan, Talofofo, and Fouha (Figure 1). Adjacent fore reef populations ∼0.5 km away were sampled to determine the extent of genetic differentiation and compare levels of genetic diversity between these two environments (Figure 1). An additional Northern fore reef population was sampled at Ritidian to determine the extent of population structure and genetic diversity on an island-wide level (Figure 1).

Massive *Porites* colonies were sampled at least 3 meters apart to avoid sampling clones. Samples were photographed and collected between 2-6 meters depth, using scuba diving and snorkeling, and 10 cm^2^ of skeleton with live tissue was removed using a hammer, chisel, and pliers. Samples were kept alive in sea water and transported back to the University of Guam (UOG) Marine Laboratory. There, half of each sample was preserved in 80% ethanol and frozen in a - 20°C freezer for genomic analysis. The remaining half was soaked in 100% bleach to remove live tissue, dried and kept as a voucher specimen for morphological analyses. Both tissue and voucher specimens for all samples are stored in the UOG Biorepository (biorepository.uog.edu). Overall, 271 massive *Porites* samples were collected for this study but only 172 were genotyped and analyzed morphologically (Table S2).

### DNA extraction, Radseq library preparation and sequencing

DNA was extracted using the Qiagen Animal Blood and Tissue DNA extraction kit (Qiagen; Hilden, Germany) and the Epoch MiniPrep GenCatch extraction kit (Epoch Life Sciences, Inc.; Missouri City, TX) following the manufacturer’s protocol. Extracted DNA was quantified using a Qubit High Sensitivity dsDNA assay (Thermo Fisher Scientific, Waltham, MA). ddRAD libraries were generated using a modified version of the double-digest RAD (ddRAD) protocols [37,38]. Briefly, extracted DNA was digested using two restriction enzymes (NsiI and MspI; New England BioLabs, Ipswich, MA). Digested DNA was then re-quantified and ligated to adaptors with sample-specific barcodes and primer annealing sites. Barcoded samples were then pooled and size-selected on a 2% agarose Invitrogen E-gel (Invitrogen, Carlsbad, CA) for fragments ranging between 350-450 bp in size. Size-selected libraries were then amplified using PCR (30 seconds at 98°C, 16-20 cycles of [10 seconds at 95°C, 30 seconds at 65°C, 60 seconds at 72°C], 5 minutes at 72°C) using pool-specific forward and reverse primers. Amplified libraries were then quantified using a Qubit High Sensitivity dsDNA assay (Thermo Fisher Scientific, Waltham, MA) and an Agilent Bioanalyzer High Sensitivity DNA assay (Agilent Technologies, Santa Clara, CA), and were then sequenced using an Illumina NextSeq500 (Illumina, Inc., San Diego, CA) at the University of Guam Marine Laboratory.

### Data curation and multi-locus genotyping

Raw sequence reads were quality-trimmed using TrimGalore 0.6.5 (https://github.com/FelixKrueger/TrimGalore) to remove low-quality reads [-q 5, --length 20, -- stringency 1, -e 0.1], and then demultiplexed using a custom-made python3 script (H. Weigand, pers. comm). After barcode removal, reads were first aligned to the *Porites lutea* bacterial metagenome and then to a *Symbiodiniaceae* C15 genome from the same *P. lutea* [39] to remove non-coral loci, using Bowtie2 [40] with default settings. Unaligned reads were then aligned to the *P*. *lutea* coral host genome [39] using Bowtie2 with default settings. Resulting bam files were then sorted and used as input for all subsequent coral genotyping and population genomic analyses.

Several different datasets were generated to make the best use of the genotype data for different analyses. To avoid linkage disequilibrium among SNPs, all population genetic analyses were conducted with datasets using only one SNP per ddRAD locus unless stated otherwise. The first dataset was a genotype likelihood dataset generated by ANGSD version 0.5 [41], containing loci present in at least 80% of all retained samples and thus called ANGSD R80. This dataset was initially used for clonality (n=97 samples, min. 100,000 reads/sample), and after clonal removal, was used for many population genetic analyses including principal coordinate (ngscovar) and admixture analyses (NGSAdmix) (n=95 samples, min. 100,000 reads/sample) since ANGSD genotype likelihoods are a more statistically robust method of genotyping for low- to medium-coverage sequence data, compared to methods utilizing genotype calls [41]. Aligned reads were converted to genotype likelihoods with ANGSD, using the following flags: -uniqueOnly 1, - remove_bads 1, -minMapQ 20, -minQ 25, -dosnpstat 1, -doHWE 1 -hetbias_pval 1e-5 - skipTriallelic 1, -snp_pval 1e-5 -minMaf 0.05 [41].

An additional dataset was generated with STACKS version **2.55** [42,43], which identifies SNPs using a were identified using a Bayesian model and makes firm genotype calls (instead of genotype likelihood scores) that are required for certain analyses [44]. Additionally, STACKS was used to produce genotype call datasets that were used to conducts several population genomic analyses in GenoDive that do not readily accept ANGSD-generated genotype likelihood data, including AMOVA and population genetic summary statistics. The main STACKS dataset contains loci present in at least 30% of all samples and is called STACKS R30. This dataset was used for phylogenetic analyses (n=172 samples, min. 4,800 reads/sample), which benefit from larger datasets, even if they are patchier. In addition, a second STACKS dataset was generated that contained loci present in at least 80% of all samples and was thus called STACKS R80. This dataset was used for AMOVA among the all clades (n=96, min. 100,000 reads/sample, Table S8), as well as among the three largest clades (n=86, n=42, Table S8). For phylogenetic and population genomic analysis, STACKS was run with default settings, except for inclusion of the cutoff to retain loci present in at least 30-80% of all samples, respectively (-r and -R flags). Additional multi-clade and clade-specific sub-datasets were also curated using both ANGSD and STACKS as outlined above and in Table S2.

### Phylogenetic and population genetic analyses

Phylogenetic analyses were based on a concatenated alignment of variable sites using the STACKS R30 dataset. The most suitable model for phylogenetic analysis was chosen by IQTree during model selection, while RAxML was run using the most widely used general time reversible model of nuclear substitution, GTRCAT. Phylogenetic analyses were conducted using a maximum-likelihood inference in RAxML 8.2.4, with model selection choosing the GTRCAT nucleotide substitution model, SNP ascertainment bias correction and 1000 bootstrap replicates on the CIPRES web portal [45]. Phylogenetic analyses were also conducted with IQTree version [46], with model selection choosing the model SYM+ASC+R4 with 1000 ultrafast bootstrap replicates and SNP ascertainment bias correction. These analyses consistently revealed the presence of several distinct clades that were treated as separate species in subsequent analyses.

Clades were assessed for their distribution across geographic locations as well as environments. Colony color was also assessed within and across clades to determine whether color is a useful proxy for in-situ clade identification.

Next, identity-by-state (IBS) pairwise relatedness matrices [47,48] were generated using ANGSD to identify clonal genotypes within each genetic clade. These matrices were then used to generate relatedness dendrograms in R [49], to provide a visualization of sample pair relatedness. Technical replicates, or multiple sequencing replicates of the same sample, were used to identify a minimum level of diversity among identical genotypes due to sequencing errors and other artifacts. Samples with relatedness values equal or greater than those of the technical replicates were identified as clones. The sample with the highest number of reads of each clonal genotype was selected as the representative sample for that genotype in subsequent analyses.

To assess and visualize patterns of genetic differentiation among samples, principal coordinate analyses were generated with the ANGSD program ngscovar [41] using the ANGSD R80 dataset. In addition, a subset of the STACKS R80 dataset was used to assess the partitioning of genetic variation among the three largest clades using AMOVA. To determine the extent of admixture among clades, admixture analyses were performed with the ANGSD subprogram NGSAdmix [41] for K values of 2 to 8 on the ANGSD R80 dataset (without the single individual representing Clade 3). To assess the validity of clades with mixed ancestry, additional admixture analyses were conducted on subsets of the larger ANGSD R80 dataset. Population genetic summary statistics across all sites (variable and invariant) were generated for observed and expected heterozygosity, as well as inbreeding coefficients, using GenoDive [50]. Additional population genetic analyses were conducted among populations within the two largest clades. An AMOVA analysis was used to assess the hierarchical partitioning of within-clade genetic diversity and pairwise FST values were generated among populations using GenoDive. Population genetic summary statistics were also calculated across all sites. For analyses of population differentiation, missing data was imputed with alleles randomly selected across populations.

### Symbiont characterization

Proportions of major Symbiodiniaceae genera were evaluated to determine if and how photosymbionts differ between clades, environments or geographic locations, as in Barfield et al. (2018). Trimmed and bacteria-filtered reads for all samples were aligned to a reference database with transcriptomes for the Symbiodinaceae genera *Breviolum, Cladocopium, Durusdinium,* and *Symbiodinium* [47,51,52] with Bowtie2 using default settings. Relative proportions of photosymbiont genera were then calculated with the custom perl script zoox.pl (https://github.com/z0on/2bRAD_denovo/). The dominant symbiont genera among samples were identified using reads with a minimum mapping quality ≥ 40.

### Morphological Assessment

To preliminarily identify characters or character sets that may be shared by clade members and to aid in the application of traditional morphology-based species determinations when such determinations were possible, qualitative assessments of several corallite-level characters were carried out, and calice diameters measured, for a haphazardly selected subset of clade member voucher specimens. Specimens were examined with the aid of a Meiji EM-51L stereo microscope and images obtained using a Nikon D7200 dSLR camera mounted to the scope. The average condition of several corallite characters was qualitatively characterized for each specimen by examining at least ten corallites; examined characters included corallite shape; degree of excavation; septal shape, thickness, and length; wall thickness; number, height, and robustness of pali; number, height and robustness of septal denticle rings; and the height, robustness, and shape of the columella. Calice diameter measurements were obtained for a haphazard selection of at least six corallites per specimen using a calibrated eyepiece reticle. Specimens known or presumed to have been obtained from the underside of the colony margin or those with a large proportion of immature corallites were excluded from the analysis.

## Results

### Reads and Loci

Across 172 sequenced samples, ddRAD sequencing yielded in 238.7 million reads across 172 samples (average =1.388 million reads/sample). After quality filtering and removing PCR duplicates, 38.36 million reads were retained (average = 258,785 reads/sample). For phylogenetic analyses, samples with ≥ 4,800 reads (n = 172, average = 223,026 reads/sample, 22,982 loci and 110,252 concatenated SNPs) were used. For population genomic analyses, only samples with ≥ 100,000 reads (n = 97, average = 360,149 reads/sample) were included.

### Phylogenetic analyses and species identity

Phylogenetic analyses with model selection using IQTree and RAxML revealed seven clearly distinct clades of massive *Porites* on Guam (Figure 2 and S1). All seven clades are significantly distinct from one another, with all but one clade (clade VI) having full node support in both phylogenetic analyses (100% bootstraps).

**Figure 2:**
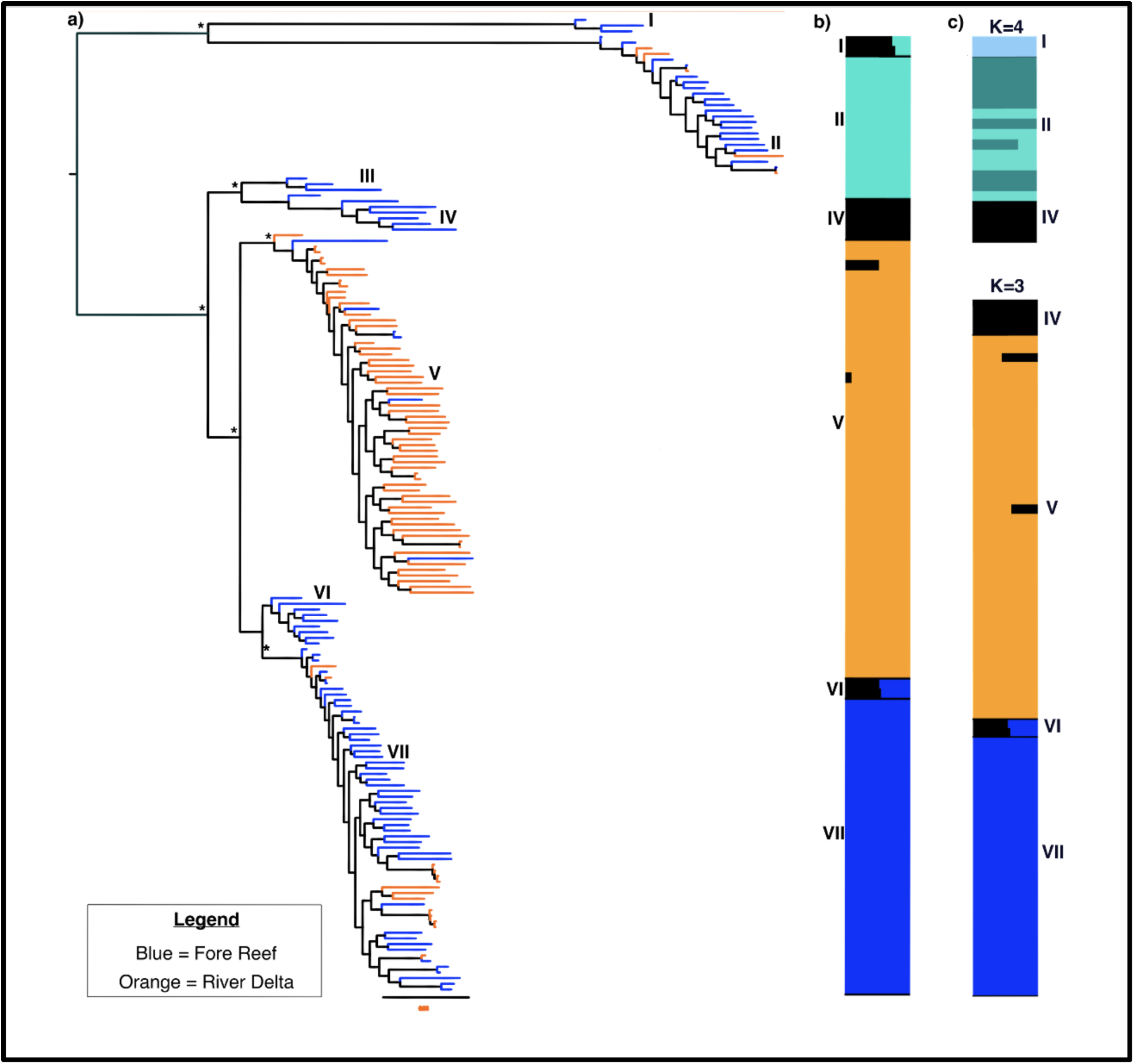
a) RAxML tree using 22,982 loci and 110,252 concatenated SNPs (lnL= -1,698,825). Node support is based on 1000 bootstraps replicates. Tip colors indicate the environmental origin of each sample. Asterisks indicate 100% bootstrap support, however, clade VI had 84% BS support. b) Admixture proportions using K=4 for clades using the ANGSD R80 dataset, which indicates a general lack of gene flow among the largest massive Porites clades. c) Top right: K=4 ANGSD R80 admixture plot including only clades I, II, and IV, indicating no ancestry of clade II and IV in clade I. Bottom right: K=3 ANGSD R30 admixture plot including only clades IV-VII, indicating clear clade IV and VII ancestry in clade VI.

The largest clade (64 specimens, 37.2% of all samples) was termed Clade V. In contrast to all other clades, it was found predominantly in river deltas (58/64 samples = 91%), representing 75% of all river delta specimens in this study (Figure 3, Table 1). The corallite morphologies of samples within this clade were fairly consistent among samples, despite the large number of samples, and are consistent with *Porites* cf. *murrayensis* (Figure 4a, Table 3) [54]. There were a few exceptions, as some samples have unusually thick septa while others have corallites with prominent pali (Table 3).

**Figure 3:**
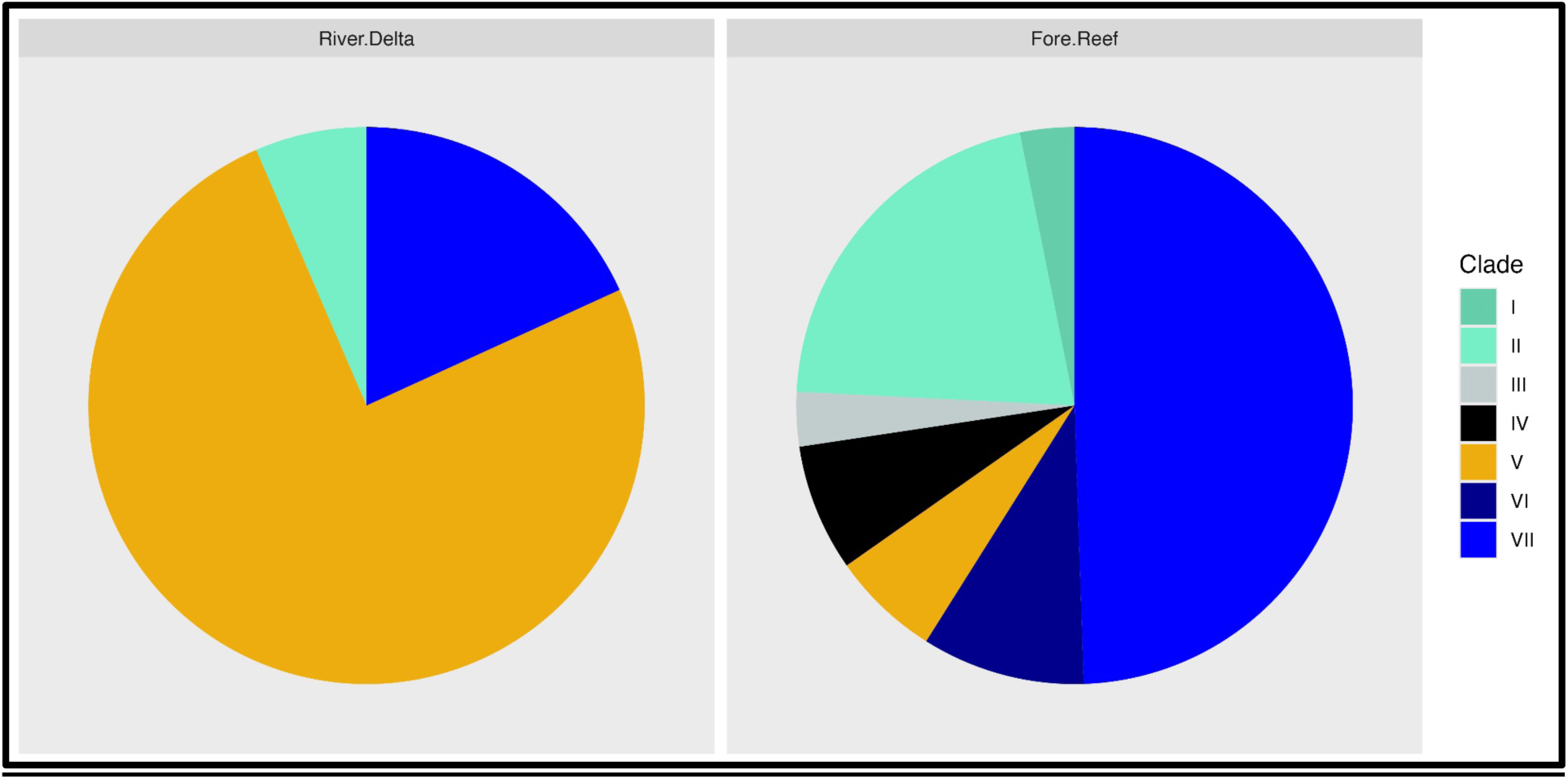
Clade representation within each habitat.

**Figure 4:**
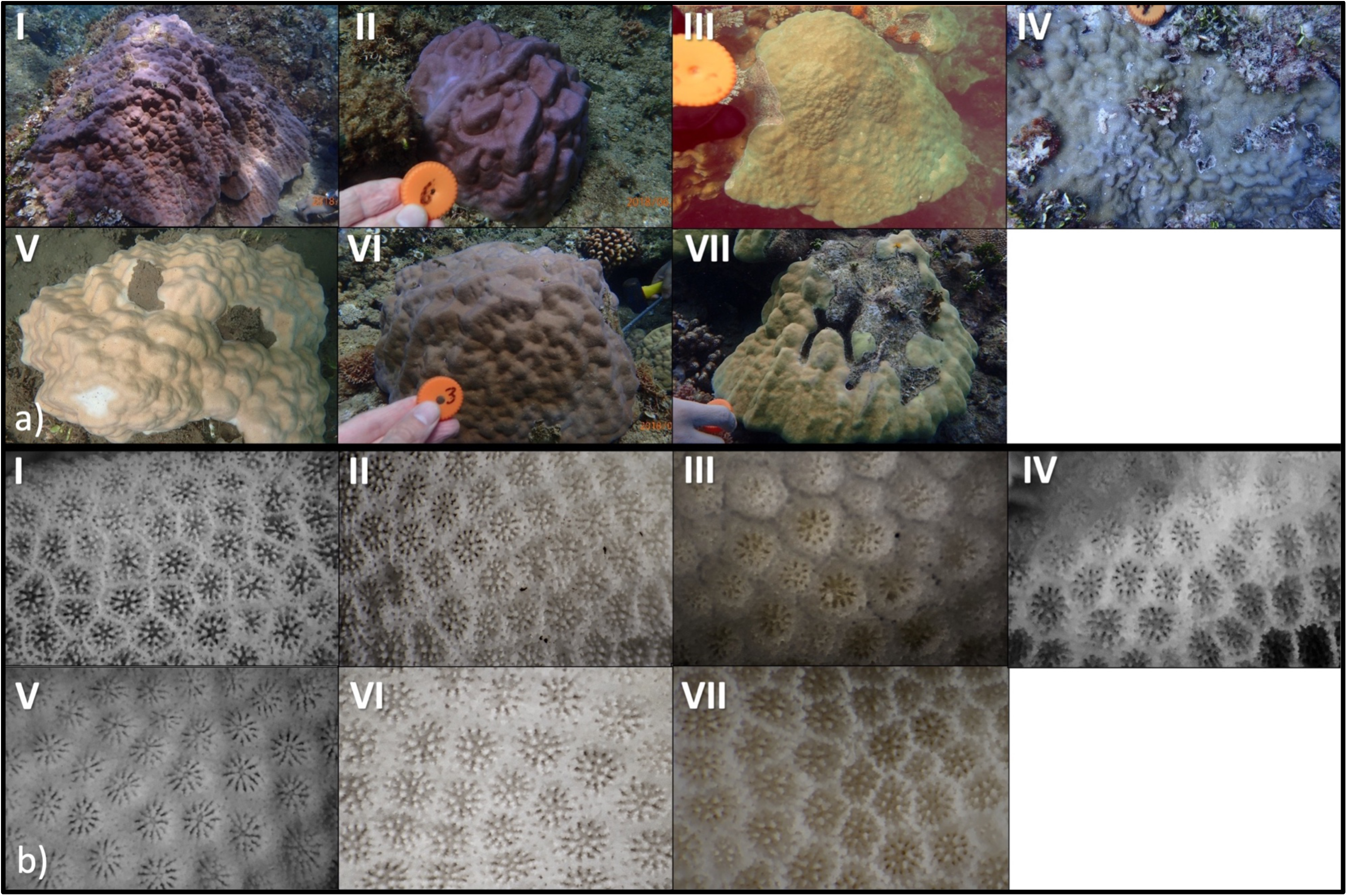
a) in-situ and b) corallite photographs of corals belonging to each of the seven clades.

**Table 1:**
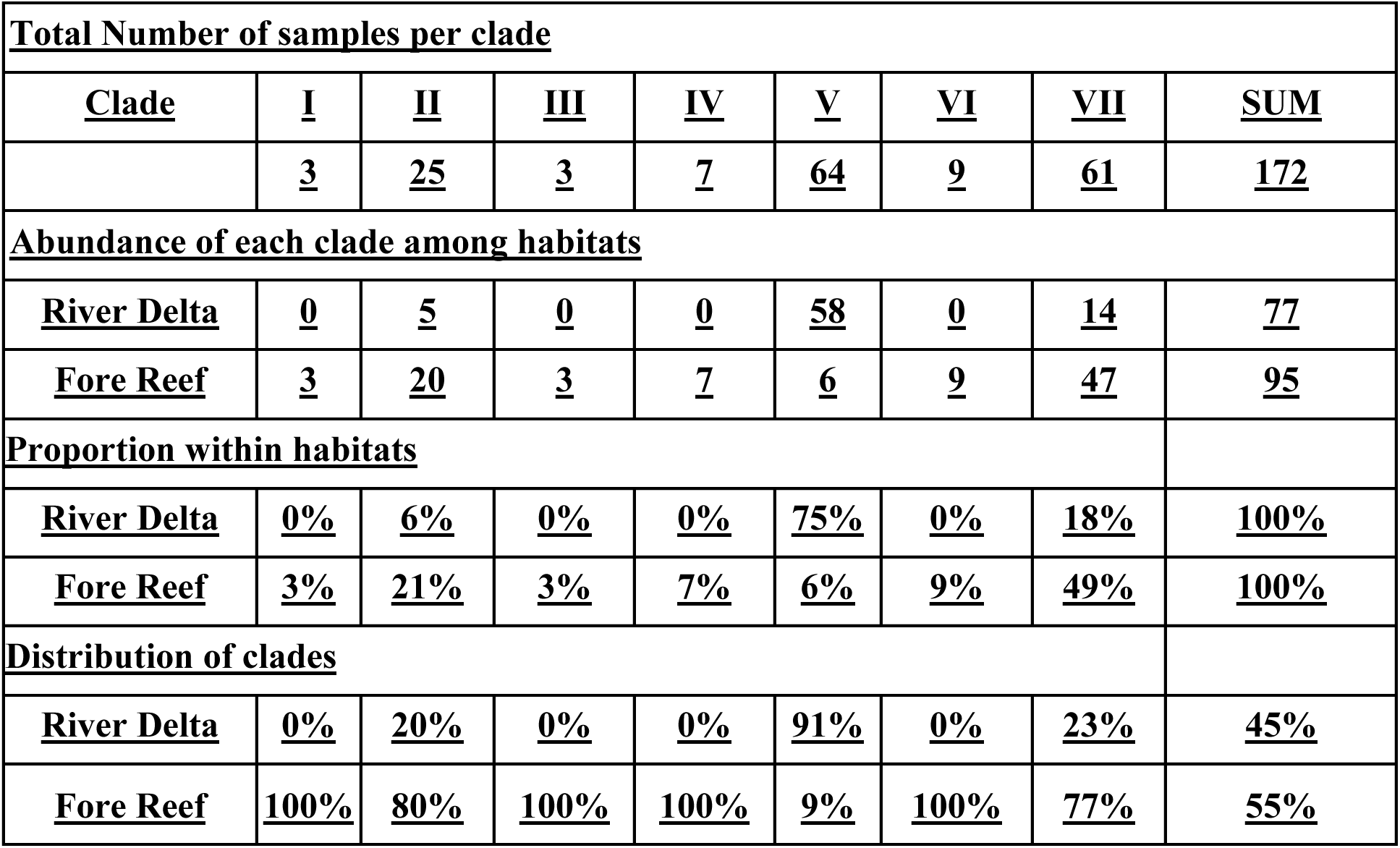
Distribution of clades over habitats.

Clade VII was the second-largest clade (61 specimens, 35.4% of all samples). It was found predominantly on fore reefs (47/61 samples; Figure 3, Table 1) and represents almost half of all fore reef specimens in this study (Table 1). Corals within this clade were identified as *Porites* cf. *australiensis* sensu Vaughan 1918 (Figure 4a) [53] but corallite morphology is more variable within this clade (Table 3).

The third-largest clade (25 samples), termed Clade II, was also found predominantly on fore reefs (20/25 samples; Figure 3, Table 1). Samples within this clade were identified morphologically as *Porites* cf*. lutea* (Figure 4a) [53].

Clade VI was found exclusively on fore reefs and 8 out of 9 samples were collected from the only Western site in this study, Fouha (Figure 3, Table 1). Samples within this clade possess corallite morphological characteristics most similar to *Porites* cf. *australiensis* (Table 3, Figure 4b) [53] as well, similar to the closely related clade VII. In the RAxML tree (Figure 2), clade VI has 84 % bootstrap support, however, in the IQTree, it has 100% bootstrap support (Figure S1).

Next, clade III (3 samples) and clade IV (7 samples) were found exclusively in fore reefs on Guam’s Eastern coast (Figure 3, Table 1). Samples within clades III and IV possess corallite morphological characteristics most similar to *Porites* cf. *lobata* [54] and *P.* cf. *solida* [55], respectively (Table 3). Differences between these clades are subtle, however, and require further investigation. All clade III specimens exhibited a mounding morphology, while all but one of clade IV specimens exhibited a thick, more encrusting colony morphology (Figure 4a).

Finally, clade I (3 samples) contains one sample each from Inarajan, Ritidian, and Fouha fore reefs (Table S4, S5). Their corallite morphology is most similar to *Porites* cf. *evermanni* (Table 3, Figure 4b) [56,57].

### Clonality

Clade-specific identity-by-state (IBS) matrices were generated in ANGSD using the ANGSD R80 dataset (Figure S2). Visual inspections of IBS cladograms for the four larger clades indicate no clones in two of these clades (IV & VII), and one pair of clones each in Clades II and V. In Clade II, both clonemates were found in Inarajan, one in the river delta and the other in the nearby fore reef (∼0.5 km apart). Interestingly, the clonal pair in Clade V was found ∼7 km apart, in the Inarajan and Talofofo river delta, respectively.

### Genetic differentiation among clades

The principal coordinate analysis (PCoA) differentiates samples into similar clusters as the phylogenetic analyses (Figure 5). Axis 1 separates the phylogenetically distinct clades I and II from the other samples, with clade I positioned between clade II and the other clades. Axis 2 separates the more closely related clades, with clades V and VII being most separated from one another and clades VI, III and IV located in between (Figure 5). Additionally, AMOVA analyses across all clades and for the three largest clades (II, V, and VII) indicates that between 61-68% of the genetic variation is partitioned among clades, providing further support for genome-wide interspecific differences between these clades (Table S8, p-value <= 0.001).

**Figure 5:**
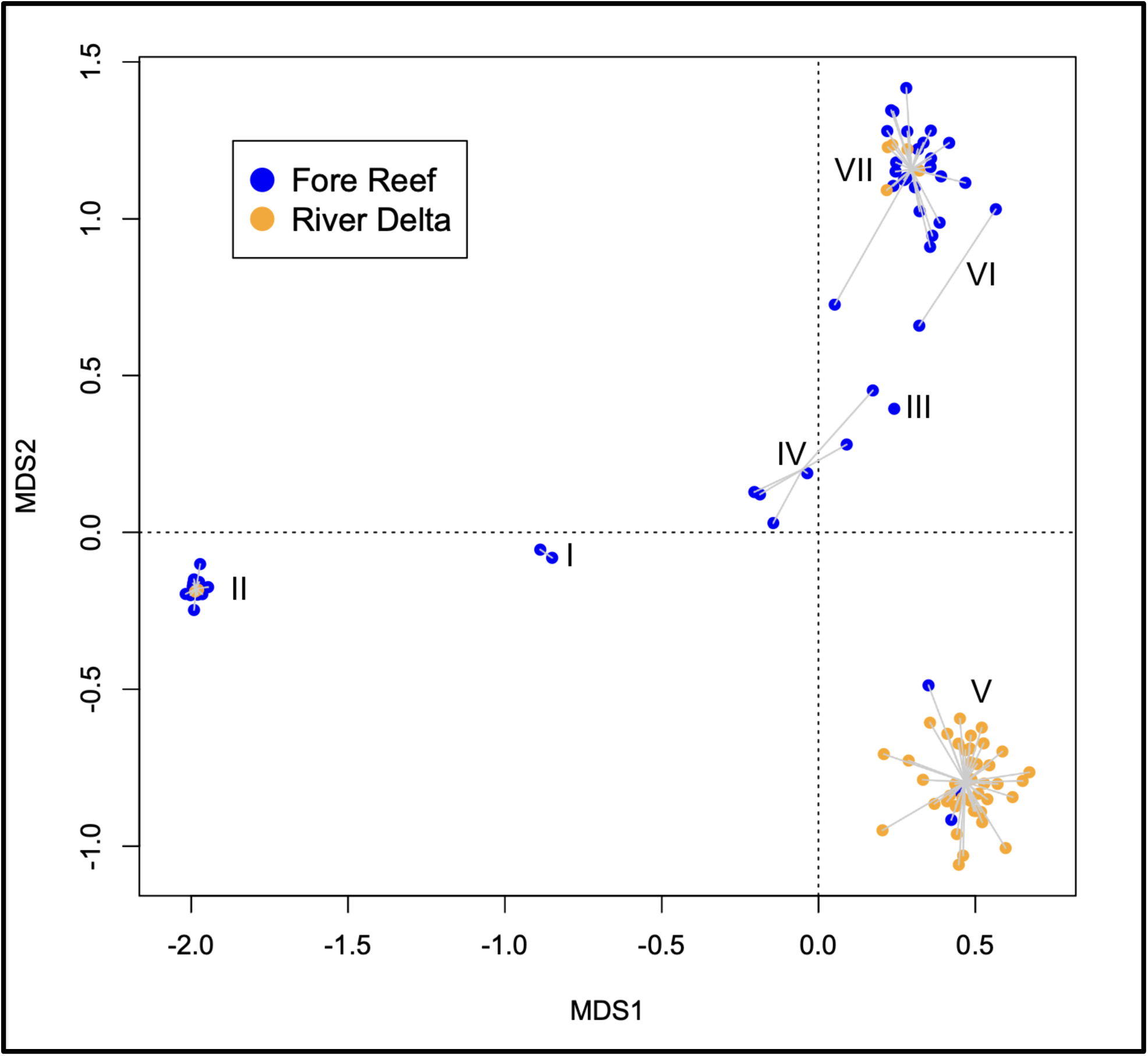
Principal coordinate analysis with all clusters corresponding to the phylogenetic clades in Figure 2. Color represents geographic location, while shape indicates the environment where each sample was collected.

Initial admixture analysis included all clades except clade III, since only one sample in this clade had more than 100,000 reads. Admixture analysis with K = 4 genetic clusters across all remaining clades show that the four main genetic clusters in this dataset (black, teal, orange, and blue) correspond respectively to the four major clades (II, IV, V, and VII; Figure 2b) with virtually no signs of admixture besides two samples in clade V (Figure 2b). These results are consistent from K=4 to K=8 (Figure S3). In contrast, the two smaller clades, Clades I and VI, showed signs of mixed ancestry. For Clade I, admixture analyses of Clades I, II, and IV (Figures 2c & S4) indicate no mixed ancestry of Clade I samples. In contrast, additional analyses for Clade VI support a mixed ancestry of this clade. Admixture analyses of clade IV-VII (Fig. S5) and IV-VI-VII (Fig. S6) consistently recover the mixed ancestry of clade VI samples (Figure 2c, S5, S6), indicating they may be hybrids of Clades IV and VII origin.

Population genetic summary statistics for the three largest clades (II, V, and VII) indicates overall comparable levels of genetic diversity (HO average = 0.0037, SD 0.004; HE = 0.0048, SD 0.0009; Table 2). Genetic diversity was slightly elevated in clade V, the most abundant clade in our dataset, respectively. Substantial inbreeding was detected in all clades (GIS average = ∼0.2) and was significantly elevated in the least abundant clade II (GIS = 0.226) compared to the two more abundant clades.

**Table 2:** Population genetic statistics for the four major clades, as well as among populations within the two largest clades (V & VII). R80S = the population genomic dataset used to calculate these statistics (see Table S2). HO = Observed Heterozygosity; He = Expected Heterozygosity; Gis = Inbreeding Coefficient; SD = Standard Deviation, 95%c.i. = 95 confidence interval. To test for differences in genetic diversity due to different sample sizes among clades, we calculated genetic diversity statistics on random 14 sample subsets of Clade V and VII (Table S8) but did not find significant differences.

| Clades/<br>Population | Samples | HO | 95% C.I. | HE | 95% C.I. | GIS | 95% C.I. |
| --- | --- | --- | --- | --- | --- | --- | --- |
| II | 14 | 0.0033 | 0.0032-0.0034 | 0.0043 | 0.0041-0.0044 | 0.226 | 0.213-0.240 |
| V | 43 | 0.0038 | 0.0037-0.0039 | 0.0046 | 0.0045-0.0046 | 0.171 | 0.155-0.171 |
| Fouha | 19 | 0.0039 |  | 0.0047 |  | 0.157 |  |
| Inarajan | 8 | 0.0038 |  | 0.0045 |  | 0.166 |  |
| Talofofo | 15 | 0.0038 |  | 0.0045 |  | 0.158 |  |
| Ritidian | 1 | - |  | - |  | - |  |
| VII | 29 | 0.0035 | 0.0034-0.0036 | 0.0042 | 0.0041-0.0043 | 0.161 | 0.157-0.174 |
| Fouha | 3 | 0.0034 |  | 0.0041 |  | 0.189 |  |
| Inarajan | 8 | 0.0036 |  | 0.0043 |  | 0.162 |  |
| Ritidian | 12 | 0.0035 |  | 0.0040 |  | 0.141 |  |
| Talofofo | 6 | 0.0036 |  | 0.0043 |  | 0.169 |  |

**Table 3:**
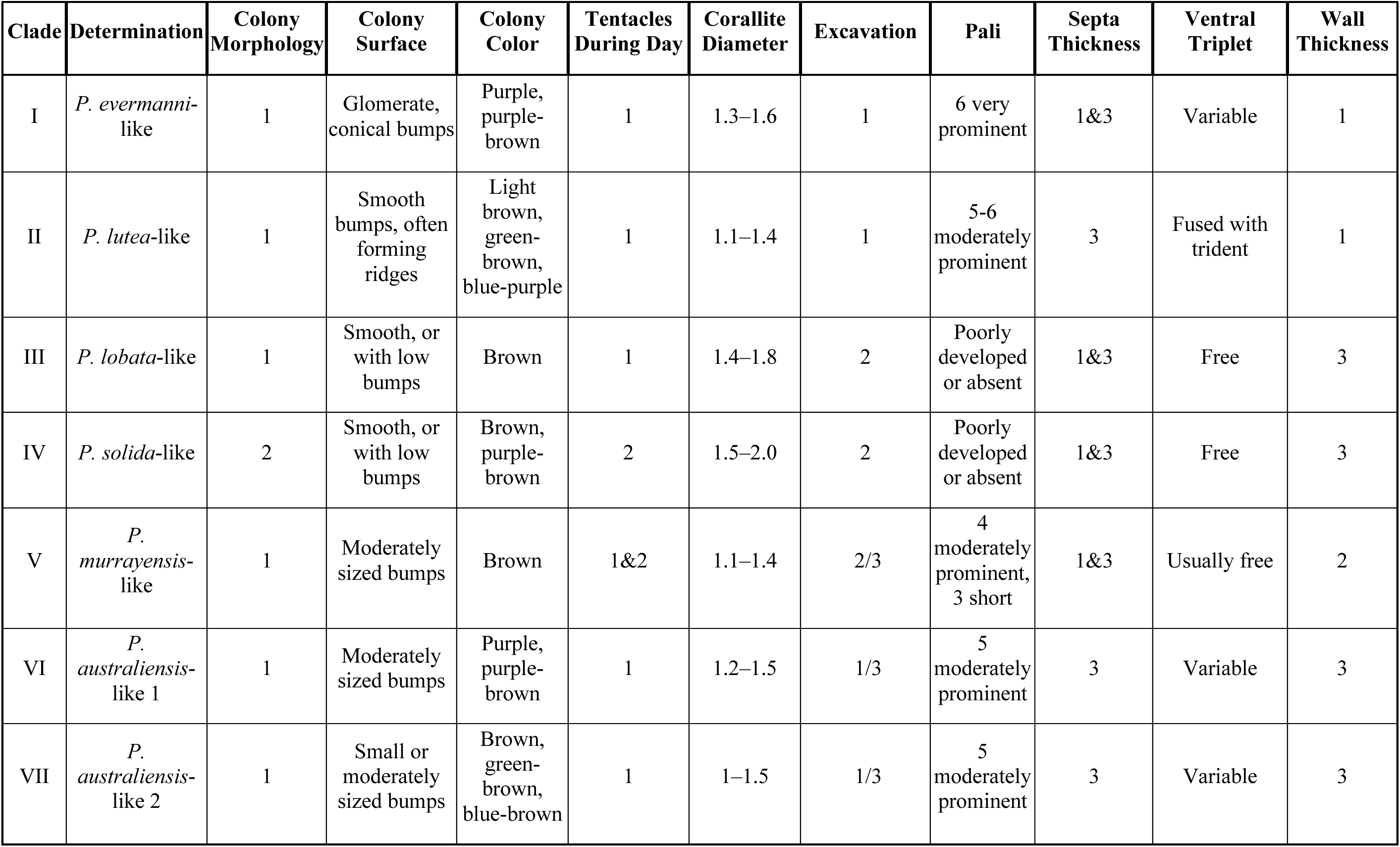
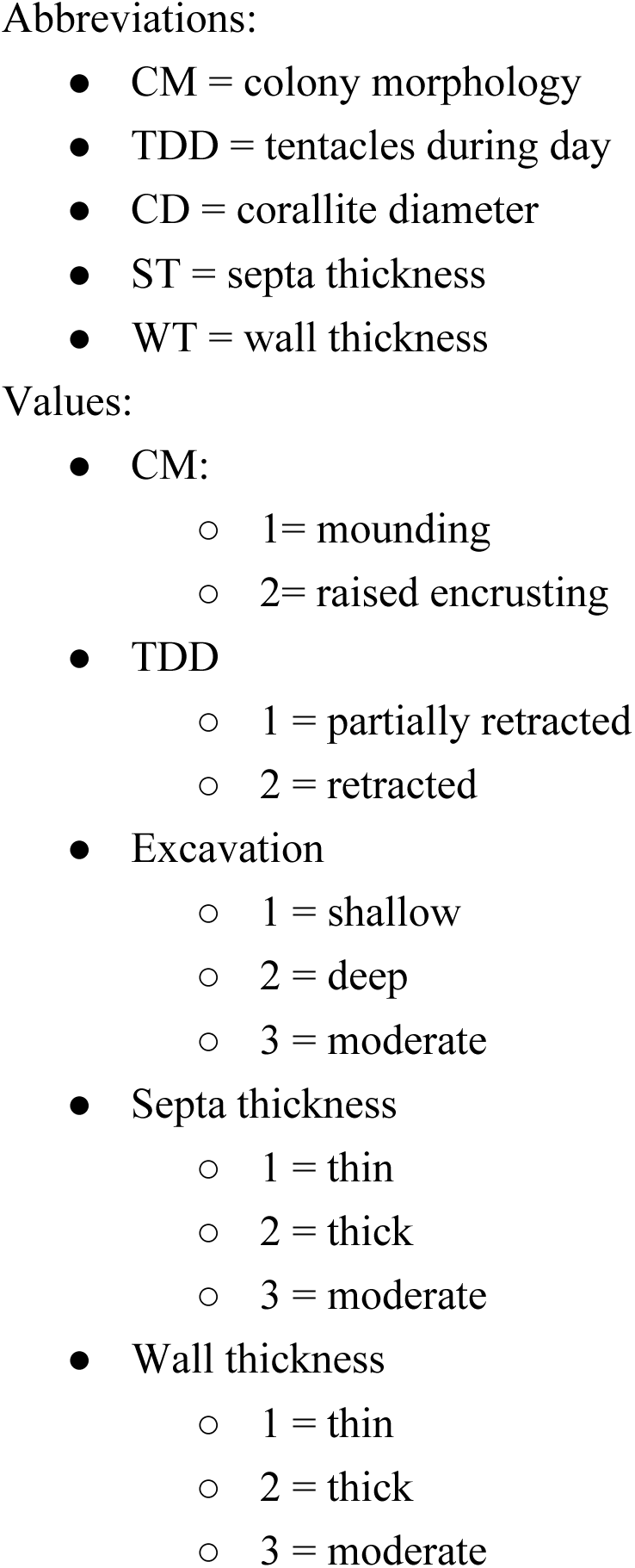
Corallite characteristics of massive Porites clades found in Guam river deltas and fore reefs.

### Population Genomic Analyses of the Two Main Clades

Only the two largest clades, Clade V (*Porites* cf*. murrayensis*) and Clade VII (*P. cf. australiensis*) had sample sizes (>15 samples) sufficient for population genomic analyses.

Clade V was predominantly found in river deltas which are concentrated in Southern Guam, yet a single clade V sample was also found on the fore reef in the far northern Ritidian population. Clade-specific AMOVA analyses, excluding the sole Ritidian sample (ORL07), indicate that 0.5% (p-value = 0.001) of its genetic diversity was partitioned among the three main populations Talofofo, Inarajan, and Fouha (Table S9). Pairwise population differentiation (*F*_ST_) was only significant for Talofofo-Fouha (P = 0.001, Table S11). This result is supported by the clade specific PCoA, which shows most differentiation between the Talofofo and Fouha populations (Figure 6a). All three populations in clade V harbored similar levels of genetic diversity, with observed heterozygosity levels ranging from 0.0038-0.0039, expected heterozygosity ranging from 0.0045-0.0047 and inbreeding coefficients ranging from 0.158-0.166 (Table 2).

**Figure 6.**
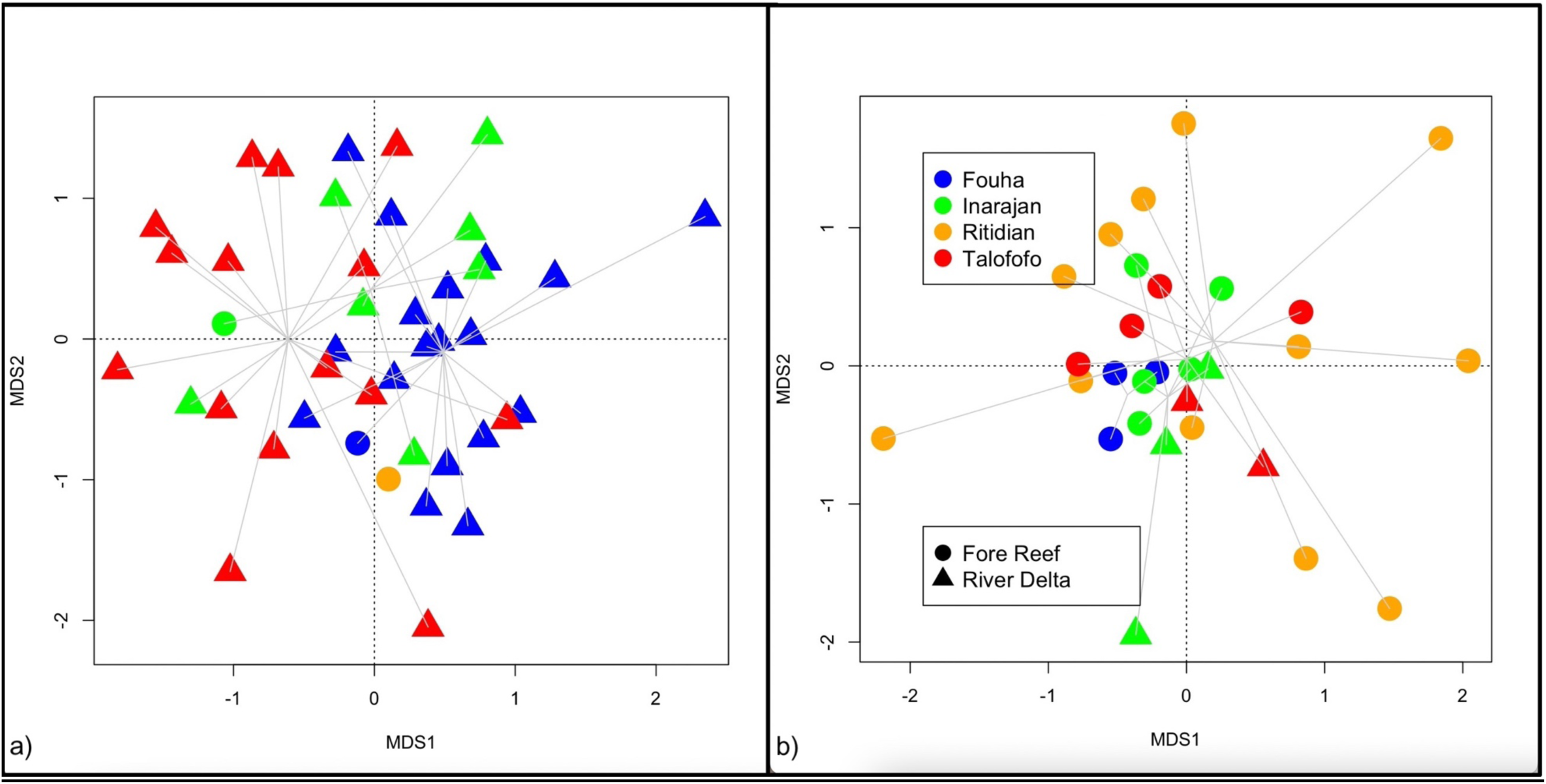
a&b: Clade V (left) and Clade VII (right) PCoAs.

Clade VII was found on all fore reef populations around Guam. Clade-specific AMOVA analyses suggest that 0.4% (p-value = 0.001) of its diversity is partitioned among the three analyzed for reef populations (excluding Fouha due to its small sample size n=3, Table S10). All pairwise *F*_ST_ values were insignificant, indicating low population structure among fore reef populations (Table S12). As expected, the clade-specific PCoA also indicates a general lack of population structure among Clade VII populations (Figure 6b). All three analyzed clade VII populations harbor similar levels of genetic diversity, with observed heterozygosity levels ranging from 0.0034-0.0036, expected heterozygosity ranging from 0.0040-0.0043 and inbreeding coefficients ranging from 0.141-0.189 (Table 2).

### Massive *Porites* Symbiont Profile Characterization

*Cladocopium* was found to be the dominant Symbiodinaceae genus in all analyzed samples, regardless of clade identity, geographic location and habitat (Figure 7). In addition, *Breviolum* and *Durusdinium* reads were detected in two samples each, but not *Symbiodinium*. Specifically, *Durusdinium* was found in clade VII in one Talofofo fore reef and one Inarajan river delta sample, while *Breviolum* was found in clade V samples from Inarajan and Talofofo’s river deltas. These results should be interpreted with caution since only 607 reads across 44 samples aligned to the symbiont reference transcriptome, averaging 13.8 reads per sample (SD = 27.3; Table S13).

**Figure 7:**
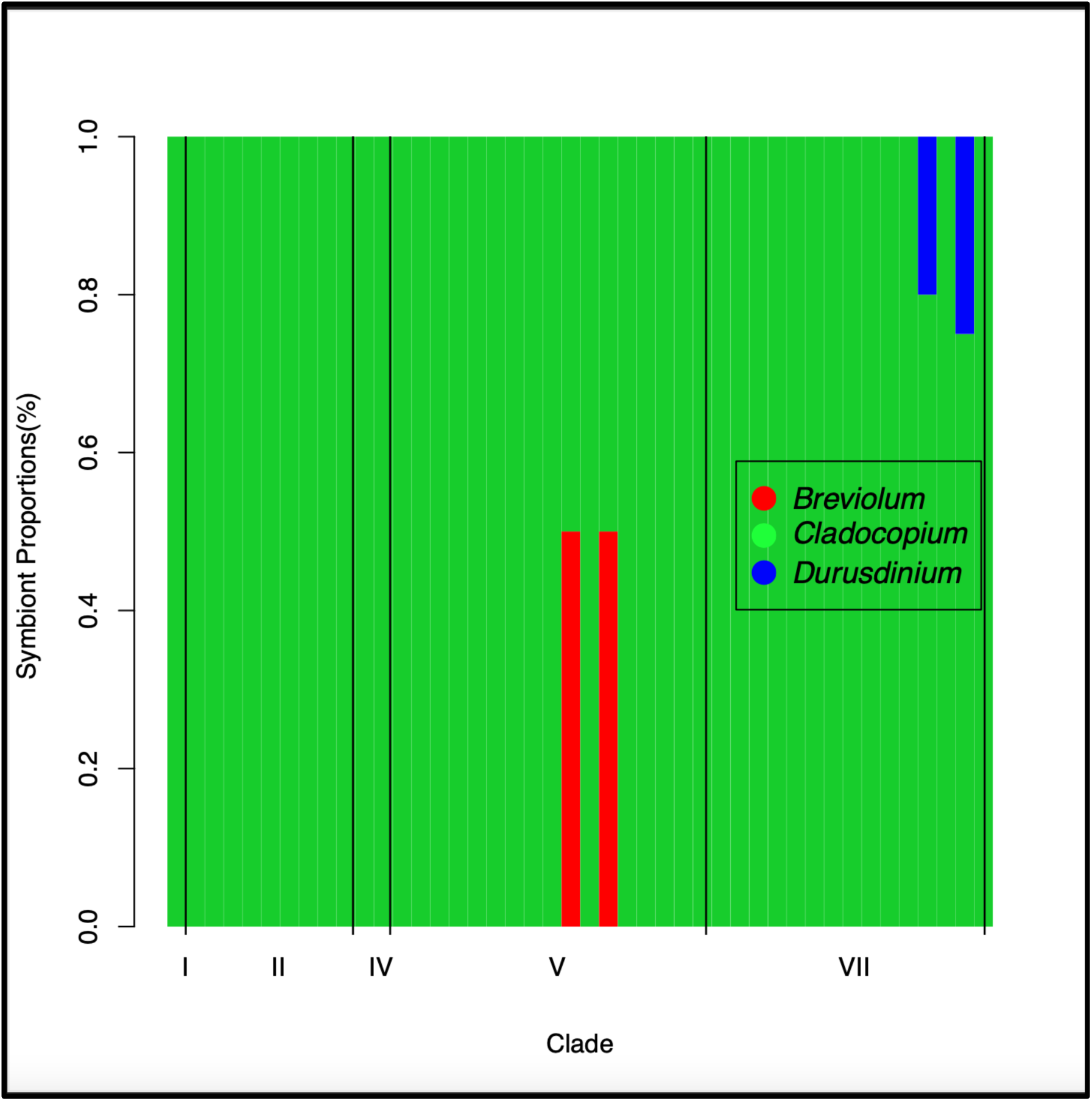
Symbiont proportions for a subset of interspecific dataset samples. Breviolum was found in one Talofofo and one Inarajan river delta sample each (RIL09 & RTL01), while Durusdinium was found in one Talofofo fore reef sample and one Inarajan river delta sample (OTL31 & RIL29).

## Discussion

### Differences in Bleaching Susceptibility Among Environments and Clades

In the current study, detailed phylogenomic and population genomic analyses of ddRADSeq data revealed that massive *Porites* in Guam’s river deltas and fore reefs are composed of seven genetically distinct but morphologically similar putative species (Figures 2, 5, S1). The abundance and distribution of the seven putative species was remarkably heterogeneous across habitats: river deltas were dominated by Clade V corals (58/77 or 75% of all river delta specimens), which were almost completely absent from fore reefs (6/95 = 6% of all fore reef specimens) (Figure 3, Table 1). In contrast, Clade II and VII corals were the most abundant species on fore reefs (67/95 specimens = 70%), while only 19 specimens across both lineages were found in river deltas (24%) (Figure 3, Table 1). The other four clades (I, III, IV, and VI) were found exclusively in fore reef habitats (Figure 3, Table 1).

Several recent studies found higher survival during bleaching events in turbid habitats compared to their offshore counterparts throughout the Indo-Pacific [1–10,13,14,58, 59–62]. This largely applies to massive *Porites* but has also been observed in numerous other genera, including *Acropora*, *Montipora*, *Turbinaria* [8], *Cyphastrea*, *Fungia* [9], *and Pocillopora* [10]. Some studies, however, specifically focus on or note differences among massive *Porites* [12–14]. Additionally, some studies even identify genetic differences between *Porites* exhibiting differential bleaching response across habitats. For example, *P. lobata* bleaches more often than the ecologically distinct yet morphologically similar massive *Porites* species *P. evermanni* [23,63]. Similarly, massive *Porites* genetic lineages in Palau’s Rock Islands have higher thermotolerance than their outer reef counterparts. In addition, cryptic, sympatric *P. lobata* lineages in Kirimati (Republic of Kiribati) have been found to exhibit differential bleaching responses and survivorship after the 2015-2016 El-Nino heatwave [64]. Additional genomic studies of *Acropora hyacinthus*, *Orbicella faveolata, and Pocillopora sp.* have identified numerous cryptic lineages that exhibit differential bleaching responses [65–68]. Together with our results, this suggests that differential bleaching responses among corals in marginal coral reefs compared to oceanic fore reefs, as well as within these habitats, may be driven by interspecific genomic differences.

Many river delta reefs are currently subject to elevated levels of pollution and runoff, which threaten the survival of corals in these marginal habitats [1,2,69,70,71]. This is especially true on Guam, where the use of off-road vehicles, wildland arson, and coastal development has accelerated the rate of terrestrial erosion and subsequent runoff [72]. Results from this study highlight that river delta habitats host genetically distinct corals compared to those from nearby oceanic corals. Therefore, efforts to minimize runoff and pollution on Guam should be undertaken to prevent biodiversity loss of these distinct marginal reef habitats. Additionally, our study highlights that habitat heterogeneity, even at local scales (<1 km), in addition to geographic distance, should be incorporated in phylogeographic and population genomic studies of reef-building corals, and more broadly in other marine taxa, to unveil the full extent of biodiversity found among and within these different habitats.

### Massive *Porites* Species and Ecological Differences

Phylogenomic analyses clearly distinguish seven distinct lineages that likely constitute seven distinct species. Admixture analyses (Fig. 2c) confirm a lack of mixed ancestry in most clades. For example, although preliminary admixture analyses indicate a mixed ancestry for Clade I, subsequent detailed analyses did not support this (Fig. S4). This is interesting since Clade I corresponds to *P. evermanni*, which has been found to hybridize with other massive *Porites* species outside of Guam [73]. One clade however, Clade VI (*P.* cf. *australiensis* I), consistently showed indications for admixture from clades IV (*P.* cf. *solida*) and VII (*P.* cf. *australiensis* II) (Fig. 2c, S4, S5). A mixed ancestry for clade VI is further supported by its low phylogenetic uniqueness, compared to other clades (Fig. 2a). Admixture between *P.* cf. *australiensis* and *P.* cf. *solida* has not been recorded before but there are numerous indications for hybridization among *Porites* species [24,73,74]. Collectively, these results suggest the presence of at least six distinct massive *Porites* species on Guam’s reefs.

These results partially align with earlier findings of multiple massive *Porites* species on Guam’s reefs based on morphological assessments. For example, *P. australiensis*, *P*. *murrayensis*, *P. lobata*, *P. lutea*, and *P. solida* were all previously identified on Guam and the Marianas Islands [33] and were delineated in the current study using genome-wide ddRAD data. Interestingly, clade I samples were morphologically identified as *P.* cf. *evermanni*, a massive *Porites* species found throughout the Central Indo-Pacific and Tropical Eastern Pacific [61], which is documented for the first time on Guam.

Our study adds to an already expansive knowledge on the ecological distribution of putative massive *Porites* morpho-species, with some species found to inhabit larger ecological ranges than previously acknowledged. For example, *P. murrayensis* (clade V) was found to be the dominant massive *Porites* coral in Guam’s river deltas, while it has been previously suggested to inhabit shallow reef flats [57,75]. *P. australiensis* and *P. lutea* (clades II & VII) were also found to occur in river deltas, while previously *P. australiensis* and *P. lutea* were thought to typically inhabit back reefs, lagoons, and fringing reefs [75,76]. On the other hand, consistent with results from the current study, *P. evermanni*, *P. lobata*, *P. solida* (clades I, III, & IV), as well as *P. australiensis* (clade VI), inhabit shallow reef environments, in addition to back reefs, lagoons, fringing reefs, and protected lagoons [75,76]. Recent studies have also highlighted ecological differences between morphologically similar massive *Porites* lineages and species. For example, distinct *P.* cf. *lobata* lineages in Palau were found to predominantly occupy either outer reef sites or the famous Rock Islands, which have consistently higher temperatures than Palau’s outer reefs [77]. In addition, morphologically similar *Porites lobata* and *P. evermanni* in the Tropical Eastern Pacific were found to predominantly inhabit either offshore or inshore habitats, respectively [23,63]. Ecological specialization, regardless of geographic distance, could therefore be a strong driver of genetic differentiation among *Porites* species and lineages.

Typically, isolation by environment and ecological differences between morphologically similar coral species have been found along depth gradients [78], with multiple cryptic lineages of putative coral species found in, yet not limited to *Montastrea cavernosa*, *Siderastrea siderea*, *Agaricia agaricites & A. fragilis*, *Pachyseris speciosa*, *Porites astreoides*, and *Eunicea flexuosa* [79–83]. Several recent studies, however, have identified ecological differences between morphologically similar species at the same depth [23,63,77,79,84]. In the current study, all species are found in at least partial sympatry on fore reefs and three species occur in sympatry in river deltas. Similar results of cryptic, sympatric lineages have recently been found for cryptic *P. lobata* lineages in Kiritimati [64], as well as *P. lobata* and *P. lutea* in Hawai’i and throughout the Indo-Pacific [17,22,85].

### Population Genomics and Phylogeography of massive *Porites*

The two largest clades in the current study, Clade V (*Porites* cf. *murrayensis*) and Clade VII (*P. cf. australiensis*), exhibited low levels of population structure around Guam (Figure 6a,6b, S7, Table S11, S12). This is not surprising considering Guam’s small geographic size (212 km²), the continuity of fore reef habitats and close proximity of Southern Guam river deltas to one another, and the fact that Indo-Pacific massive *Porites* are broadcast spawners [22,73, 86–90].

Genetic diversity, specifically observed and expected heterozygosity, in clades V and VII was broadly similar. Clade VII harbored higher levels of heterozygosity than clade V (Table 1), which is likely due to clade VII being more abundant both on Guam and likely throughout Micronesia than clade V, which on Guam, is largely confined to river deltas. Interestingly, Guam has by far the largest number of rivers among all other Marianas islands and across Micronesia (Digital Atlas of Micronesia), only comparable to Palau. This highlights the importance of Guam’s river delta habitats for the survival of massive *Porites* species diversity in the greater Micronesian region.

Genetic diversity estimates for clade V and VII populations (Table 2) fall within the lower range of genetic diversity estimates of other Indo-Pacific *Porites,* as well as other corals and other marine organisms on Guam [13,86,91–95]. Micronesian reefs are experiencing intense thermal stress and frequent bleaching events [92]. In addition, Guam, similar to the Eastern Tropical Pacific, is a largely isolated reef system, lying in the North Equatorial Current (NEC), which flows westward, with minimal larval inflow as no islands are found upstream or downstream for ∼2,000 km [96,97]. Guam massive *Porites* may therefore experience future declines in genetic diversity and the resilience needed to withstand climate change stressors.

### Symbionts of Massive *Porites* lineages among and within habitats on Guam

*Cladocopium* was found to be the dominant symbiont genus among all massive *Porites*, irrespective of clade identity, location or habitat (Figure 7). Symbiont community composition among river delta and fore reef massive *Porites* on Guam may therefore not contribute to or drive ecological specialization of massive *Porites* in these starkly distinct habitats. This is not surprising since *Cladocopium* is the most common coral symbiont throughout the Indo-Pacific and Caribbean [98,99]. Likewise, among massive *Porites* corals, *Cladocopium* was found to be the major algal symbiont among *P. lutea and P. lobata* in Singapore, Malaysia, Hawai’i, Kirimati, Palau, and the broader Eastern Tropical Pacific [85,100–102].

Ecological specialization has in some cases been attributed to symbiont coevolution in corals and other cnidarians [103]. For example, throughout the Caribbean, *Eunicea flexuosa* cryptic lineages which are largely segregated by depth host distinct symbiont phylotypes [104]. Additionally, cryptic lineages of *Madracis pharensis* segregated by depth also host distinct symbiont profiles [105]. Moreover, ecologically specialized massive *Porites* in Kirimati, which exhibited bleaching variation and differential mortality hosted distinct *Cladocopium* symbiont communities [64]. In some cases, however, symbiont communities may not differ among habitat-specialized coral species. For example, *P. lobata* and *P. compressa*, two ecologically and morphologically distinct *Porites* species that undergo high gene flow in Hawai’i, exhibited no differences in symbiont community profiles [24].

It still remains unclear, however, both whether and to what extent ecologically significant variation may be present between *Cladocopium* communities in river delta and fore reef massive *Porites* on Guam. The low number of reads, likely due to our coral host-targeted genotyping approach, limits the analytical depth and interpretability of our results. The genus *Cladocopium* is notably diverse, likely containing hundreds of species [98,99]. An ever-expanding number of formally recognized species have been discovered even as recently as 2023 [99]. Future work should employ symbiont-targeted genetic markers, such as the large ribosomal subunit (LSU), *psb*A minicircle non-coding region (*psb*A), mitochondrial cytochrome oxidase I (*coxI*), the internal transcribed spacer II (*ITS2*) region, and microsatellites to resolve any ecologically significant genetic substructuring of *Cladocopium* among Guam fore reef and river delta massive *Porites* [106–109].

## Conclusion

In this study, we identified multiple species of massive *Porites* on Guam in river deltas and adjacent fore reefs, with each of these habitats being dominated by a different clade. Differential bleaching response between these habitats may therefore be driven by interspecific differences between coral hosts rather than their symbionts, in addition to environmental differences. Despite evidence suggesting that marginal coral reefs could serve as potential refuges for corals during bleaching events, genome-wide differences between corals in marginal reefs and nearby oceanic fore reefs suggest that marginal coral reefs may only provide species-specific refuge during bleaching events.

*Porites*, and especially massive *Porites*, is in need of further taxonomic revision given the extent of phenotypic plasticity and morphological similarity within the genus, as well as its abundance and role as a major reef-building coral genus. Therefore, integrated genomic and micromorphological work of *Porites* is needed to gain a clear understanding of the extent of *Porites* diversity, both within and among different environments. For example, new barcoding markers [34,110] are extremely useful to clarify the ecological preferences and distribution ranges of cryptic species. Additionally, it is crucial to identify whether population or species-level genetic differences are found between other morphologically similar, ecologically significant reef-building corals in other habitats on Guam’s reefs, such as back reefs, lagoons, and reef crests, as well as between starkly different, adjacent habitats in other Micronesian islands, and more broadly in other Indo-Pacific islands.

## Acknowledgement

We would like to thank Dareon Rios for laboratory, bioinformatic, and analytical assistance. We would also like to thank Jessica Fernandez and Khanh Ly for laboratory assistance. We would also like to thank Hannah Weigand for providing the demultiplexing python3 script, as well as Andrew McInnis, Constance Sartor, Ka’ohinani Kawahigashi, Mariel Cruz, John Peralta, Jason Miller, and Joseph Cummings for assisting with sample collections. We would also like to thank Alexa Huzar and Carly Scott for manuscript edits. This material is based upon work supported by the University of Guam and the National Science Foundation under grant number OIA:1457769.

## Author Contributions

Conceptualization: D.C.

Methodology: D.C., K.P.

Writing-review and editing: K.P., D.C., D.B., S.L, Z.F

Formal analysis and investigation: K.P., D.C., D.B.

Data curation, visualization, and writing:

Original draft preparation: Lead: K.P, Supporting: D.C., D.B., S.L, Z.F

Funding acquisition, project administration, and resources: D.C.

## Data Availability Statement

Raw fastq files for all 172 samples are available on the NCBI Short Read Archive (SRA) under BioProject number PRJNA1107942. Scripts for bioinformatic and data analysis are available on Github at https://github.com/karimprimov/guam_massive_porites.git.

## Additional Information

### Competing Interests

The authors declare no competing interests.

## Notes

### Competing Interest Statement

The authors have declared no competing interest.

