## Supplementary Material for "Genomic data reveals habitat partitioning in massive *Porites* on Guam, Micronesia"

**Table S1:** GPS Coordinates of Guam massive *Porites* sampling locations.

| Location | Environment | GPS Coordinate |
|---|---|---|
| Inarajan | Fore reef | 13°16.524' N, 144°45.430' E |
| Inarajan | River Delta | 13°16.6169' N, 144°45.0357' E |
| Talofofo | Fore reef | 13°20.685' N, 144°46.469' E |
| Talofofo | River Delta | 13°20.22' N, 144°46.0817' E |
| Fouha | Fore reef | 13°18.45' N, 144°39.20' E |
| Fouha | River Delta | 13°18.3637' N, 144°39.416' E |
| Ritidian | Fore reef | 13°37.903' N, 144°53.677' E |

**Table S2:** Clade-specific sample read count chart with number of samples included in different datasets, as well as Specify catalog numbers in the University of Guam Biorepository (biorepository.uog.edu).

**Legend:**

Populations:

OFL=Fouha fore reef
RFL= Fouha river delta
OIL= Inarajan fore reef
RIL= Inarajan river delta
OTL= Talofofo fore reef
RTL= Talofofo river delta
ORL= Ritidian fore reef

**Notes:**

**Coral filtered reads**: Symbiont and bacteria-free reads that align to the *P.lutea* coral host reference genome.

**Phylogenetic R30:** Most comprehensive dataset that includes all samples with at least 4800 coral reads (n = 172). This dataset was used only for phylogenetic analyses (Fig 2a & Fig. S1).

**Popgen R80A:** Basic population genetic dataset, including all samples with at least 100,000 reads. This dataset was used for clonality analysis (n = 97) for clades II, IV, V, and VII (Fig. S2) with ANGSD. In addition, this dataset was used for ANGSD-based PCoA (Fig. 5) and Admixture (Fig. 2b&c) after removing two clonal samples (OIL11 from Clade II and RIL28 from clade V).

**Popgen R80S:** This dataset was used for Stacks-based Population Genetic Statistics (Table 2), AMOVAs (Table S9) and $F_{ST}$ (Table S12 & S13), which focussed on the three largest clades (II, V and VII) and excluded both clonal samples (n = 86).

**Clade-specific Popgen R80A & R80S:** Datasets used for clade-specific population genomic analyses comparing genetic diversity and connectivity among populations within clades V & VII (Tables 2, S12 & S13, Figures 6 & S7)

Clade I

| Sample | Catalog # | Coral Reads | NCBI Accession number | Dataset presence/absence | | |
|---|---|---|---|---|---|---|
| | | | | Phylogenetic R30 | Popgen R80A | Popgen R80S |
| | | | | 3 | 2 | 0 |
| OFL07 | 56 | 372,287 | SAMN41218138 | X | X | |
| OIL25 | 1065 | 200,663 | SAMN41218296 | X | X | |

| | | | | |
|---|---|---|---|---|
| ORL17 | 965 | 73,506 | SAMN41218244 | X |

Clade I was too small to be included in subsequent population genetic analyses with the R80S dataset.

Clade II

| Sample | Catalog # | Coral Reads | NCBI Accession number | Dataset presence/absence | | |
|---|---|---|---|---|---|---|
| | | | | Phylogenetic R30 | Popgen R80A | Popgen R80S |
| | | | | 25 | 15 | 14 |
| OFL04 | 53 | 420,966 | SAMN41218135 | X | X | X |
| OFL06 | 55 | 355,664 | SAMN41218137 | X | X | X |
| OFL08 | 57 | 270,974 | SAMN41218139 | X | X | X |
| OFL30 | 79 | 107,969 | SAMN41218146 | X | X | X |
| OIL10 | 1050 | 23,429 | SAMN41218285 | X | | |
| OIL11 | 1051 | 177,629 | SAMN41218286 | X | [clone][1] | - |
| OIL14 | 1054 | 606,983 | SAMN41218289 | X | X | X |
| OIL19 | 1059 | 331,454 | SAMN41218291 | X | X | X |
| OIL21 | 1061 | 542,067 | SAMN41218293 | X | X | X |
| OIL22 | 1062 | 338,368 | SAMN41218294 | X | X | X |
| OIL26 | 1066 | 343,569 | SAMN41218297 | X | X | X |
| OIL29 | 1069 | 93,811 | SAMN41218299 | X | | |
| OIL32 | 72 | 430,156 | SAMN41218300 | X | X | X |
| ORL01 | 949 | 74,903 | SAMN41218229 | X | | |
| ORL02 | 950 | 12,614 | SAMN41218230 | X | | |
| ORL31 | 979 | 73,781 | SAMN41218251 | X | | |
| ORL38 | 986 | 87,496 | SAMN41218253 | X | | |

| | | | | | | |
|---|---|---|---|---|---|---|
| OTL21 | 1025 | 446,167 | SAMN41218269 | X | X | X |
| OTL35 | 1039 | 58,642 | SAMN41218278 | X | | |
| OTL36 | 1040 | 255,223 | SAMN41218279 | X | X | X |
| RFL16 | 105 | 53,607 | SAMN41218159 | X | | |
| RFL29 | 118 | 1,331,143 | SAMN41218172 | X | X | X |
| RIL10 | 452 | 78,103 | SAMN41218187 | X | | |
| RIL11 | 453 | 1,103,269 | SAMN41218188 | X | X | X[1] |
| RTL13 | 495 | 83,280 | SAMN41218214 | X | | |

**Note:** [1] Clonality analyses with the Popgen R80A dataset indicated that OIL11 & RIL11 are clones. OIL11 was therefore excluded from the Popgen R80S dataset and all population genetic analyses.

Clade III

| Sample | Catalog # | Coral Reads | NCBI Accession number | Dataset presence/absence | | |
|---|---|---|---|---|---|---|
| | | | | Phylogenetic R30 | Popgen R80A | Popgen R80S |
| | | | | 3 | 1 | 0 |
| OIL35 | 1075 | 462,020 | SAMN41218302 | X | X | |
| ORL39 | 987 | 23,565 | SAMN41218254 | X | | |
| OTL26 | 1030 | 55,122 | SAMN41218272 | X | | |

<u>Clade III was too small to be included in subsequent population genetic analyses with the R80S dataset.</u>

Clade IV

| Sample | Catalog # | Coral Reads | NCBI Accession number | Dataset presence/absence | | |
|---|---|---|---|---|---|---|
| | | | | Phylogenetic R30 | Popgen R80A | Popgen R80S |
| | | | | 7 | 6 | 0 |

| | | | | | |
|---|---|---|---|---|---|
| OIL15 | 1055 | 128,367 | SAMN41218290 | X | X |
| OIL28 | 1068 | 150,266 | SAMN41218298 | X | X |
| ORL19 | 967 | 28,146 | SAMN41218246 | X | |
| OTL10 | 1015 | 103,897 | SAMN41218264 | X | X |
| OTL19 | 1023 | 709,611 | SAMN41218267 | X | X |
| OTL20 | 1024 | 101,086 | SAMN41218268 | X | X |
| OTL32 | 1036 | 328,953 | SAMN41218276 | X | X |

Clade V was too small to be included in subsequent population genetic analyses with the R80S dataset.

Clade V

| Sample | Catalog # | Coral Reads | NCBI Accession number | Dataset presence/absence | | | | |
|---|---|---|---|---|---|---|---|---|
| | | | | **Phylogenetic R30** | **Popgen R80A** | **Popgen R80S** | **Clade V-R80A** | **Clade V-R80S** |
| | | | | 64 | 44 | 43 | 43 | 43 |
| OFL22 | 71 | 71,380 | SAMN41218142 | X | | | | |
| OFL36 | 85 | 1,335,322 | SAMN41218150 | X | X | X | X | X |
| OIL04 | 1044 | 75,994 | SAMN41218282 | X | | | | |
| OIL12 | 1052 | 118,671 | SAMN41218287 | X | X | X | X | X |
| OIL13 | 1053 | 7,334 | SAMN41218288 | X | | | | |
| ORL07 | 955 | 217,688 | SAMN41218235 | X | X | X[1] | [1] | [1] |
| RFL04 | 93 | 855,087 | SAMN41218152 | X | X | X | X | X |
| RFL07 | 96 | 246,862 | SAMN41218153 | X | X | X | X | X |
| RFL08 | 97 | 502,179 | SAMN41218154 | X | X | X | X | X |
| RFL09 | 98 | 292,725 | SAMN41218155 | X | X | X | X | X |

| | | | | | | | | |
|---|---|---|---|---|---|---|---|---|
| RFL11 | 100 | 9,783 | SAMN41218156 | X | | | | |
| RFL14 | 103 | 496,450 | SAMN41218157 | X | X | X | X | X |
| RFL15 | 104 | 719,516 | SAMN41218158 | X | X | X | X | X |
| RFL17 | 106 | 5,571 | SAMN41218160 | X | | | | |
| RFL18 | 107 | 111,895 | SAMN41218161 | X | X | X | X | X |
| RFL19 | 108 | 4,842 | SAMN41218162 | X | | | | |
| RFL20 | 109 | 45,904 | SAMN41218163 | X | | | | |
| RFL21 | 110 | 75,535 | SAMN41218164 | X | | | | |
| RFL22 | 111 | 292,936 | SAMN41218165 | X | X | X | X | X |
| RFL23 | 112 | 110,510 | SAMN41218166 | X | X | X | X | X |
| RFL24 | 113 | 54,364 | SAMN41218167 | X | | | | |
| RFL25 | 114 | 154,641 | SAMN41218168 | X | X | X | X | X |
| RFL26 | 115 | 1,047,948 | SAMN41218169 | X | X | X | X | X |
| RFL27 | 116 | 22,541 | SAMN41218170 | X | | | | |
| RFL28 | 117 | 21,389 | SAMN41218171 | X | | | | |
| RFL30 | 119 | 218,431 | SAMN41218173 | X | X | X | X | X |
| RFL31 | 120 | 1,126,680 | SAMN41218174 | X | X | X | X | X |
| RFL32 | 121 | 339,532 | SAMN41218175 | X | X | X | X | X |
| RFL33 | 122 | 1,124,169 | SAMN41218176 | X | X | X | X | X |
| RFL34 | 123 | 222,256 | SAMN41218177 | X | X | X | X | X |
| RFL36 | 125 | 1,438,881 | SAMN41218178 | X | X | X | X | X |
| RFL37 | 126 | 204,900 | SAMN41218179 | X | X | X | X | X |
| RFL39 | 128 | 63,023 | SAMN41218180 | X | | | | |
| RIL01 | 443 | 297,757 | SAMN41218181 | X | X | X | X | X |

| | | | | | | | | |
|---|---|---|---|---|---|---|---|---|
| RIL02 | 444 | 26,336 | SAMN41218182 | X | | | | |
| RIL07 | 449 | 186,329 | SAMN41218184 | X | X | X | X | X |
| RIL08 | 450 | 87,267 | SAMN41218185 | X | | | | |
| RIL09 | 451 | 329,019 | SAMN41218186 | X | X | X | X | X |
| RIL12 | 454 | 335,458 | SAMN41218189 | X | X | X | X | X |
| RIL14 | 456 | 41,166 | SAMN41218190 | X | | | | |
| RIL15 | 457 | 125,506 | SAMN41218191 | X | X | X | X | X |
| RIL17 | 459 | 235,436 | SAMN41218192 | X | X | X | X | X |
| RIL24 | 466 | 120,003 | SAMN41218198 | X | X | X | X | X |
| RIL28 | 470 | 129,045 | SAMN41218200 | X | [clone][2] | - | - | - |
| RIL36 | 478 | 10,118 | SAMN41218205 | X | | | | |
| RIL38 | 480 | 60,664 | SAMN41218206 | X | | | | |
| RTL01 | 483 | 333,227 | SAMN41218207 | X | X | X | X | X |
| RTL08 | 490 | 152,326 | SAMN41218209 | X | X | X | X | X |
| RTL09 | 491 | 177,321 | SAMN41218210 | X | X | X | X | X |
| RTL10 | 492 | 416,669 | SAMN41218211 | X | X | X | X | X |
| RTL11 | 493 | 276,894 | SAMN41218212 | X | X | X | X | X |
| RTL12 | 494 | 243,820 | SAMN41218213 | X | X | X | X | X |
| RTL15 | 497 | 203,782 | SAMN41218215 | X | X | X | X | X |
| RTL16 | 498 | 452,346 | SAMN41218216 | X | X | X | X | X |
| RTL17 | 499 | 113,076 | SAMN41218217 | X | X | X | X | X |
| RTL18 | 500 | 523,079 | SAMN41218218 | X | X | X | X | X |
| RTL20 | 502 | 145,480 | SAMN41218219 | X | X | X | X | X |
| RTL21 | 503 | 5,530 | SAMN41218220 | X | | | | |

| | | | | | | | | |
|---|---|---|---|---|---|---|---|---|
| RTL22 | 504 | 13,003 | SAMN41218221 | X | | | | |
| RTL23 | 505 | 70,926 | SAMN41218222 | X | | | | |
| RTL24 | 506 | 128,068 | SAMN41218223 | X | X | X | X | X |
| RTL25 | 507 | 141,620 | SAMN41218224 | X | X | X | X | X |
| RTL26 | 508 | 258,730 | SAMN41218225 | X | X | X | X | X |
| RTL33 | 515 | 130,170 | SAMN41218226 | X | X | X | X | X |

**Note:** [1] ORL07 was the only sample from Ritidian and was thus excluded from population geomic analyses.
[2] Clonality analyses with the Popgen R80A dataset indicated that RIL28 & RTL25 are clones. RIL28 was therefore excluded from the Popgen R80S dataset and all population genetic analyses.

Clade VI

| Sample | Catalog # | Coral Reads | NCBI Accession number | Dataset presence/absence | | |
|---|---|---|---|---|---|---|
| | | | | Phylogenetic R30 | Popgen R80A | Popgen R80S |
| | | | | 9 | 2 | 0 |
| OFL01 | 50 | 7,063 | SAMN41218132 | X | | |
| OFL02 | 51 | 41,105 | SAMN41218133 | X | | |
| OFL03 | 52 | 29,337 | SAMN41218134 | X | | |
| OFL05 | 54 | 14,132 | SAMN41218136 | X | | |
| OFL23 | 72 | 153,227 | SAMN41218143 | X | X | - |
| OFL27 | 76 | 40,159 | SAMN41218145 | X | | |
| OFL34 | 83 | 70,896 | SAMN41218149 | X | | |
| OFL38 | 87 | 56,237 | SAMN41218151 | X | | |
| OTL24 | 1028 | 104,493 | SAMN41218271 | X | X | - |

Clade VII

| Sample | Catalog # | Coral Reads | NCBI Accession number | Dataset presence/absence | | | | |
|---|---|---|---|---|---|---|---|---|
| | | | | **Phylogenetic R30** | **Popgen R80A** | **Popgen R80S** | **CladeVII-R80A** | **CladeVII-R80S** |
| | | | | 61 | 29 | 29 | 29 | 29 |
| OFL12 | 61 | 44,939 | SAMN41218140 | X | | | | |
| OFL18 | 67 | 682,981 | SAMN41218141 | X | X | X | X | X |
| OFL24 | 73 | 60,456 | SAMN41218144 | X | | | | |
| OFL32 | 81 | 115,311 | SAMN41218147 | X | X | X | X | X |
| OFL33 | 82 | 229,681 | SAMN41218148 | X | X | X | X | X |
| OIL01 | 1041 | 74,213 | SAMN41218280 | X | | | | |
| OIL03 | 1043 | 159,058 | SAMN41218281 | X | X | X | X | X |
| OIL05 | 1045 | 135,826 | SAMN41218283 | X | X | X | X | X |
| OIL09 | 1049 | 5,647 | SAMN41218284 | X | | | | |
| OIL20 | 1060 | 126,706 | SAMN41218292 | X | X | X | X | X |
| OIL24 | 1064 | 11,717 | SAMN41218295 | X | | | | |
| OIL33 | 1073 | 325,234 | SAMN41218301 | X | X | X | X | X |
| OIL40 | 1080 | 354,808 | SAMN41218303 | X | X | X | X | X |
| ORL03 | 951 | 66,047 | SAMN41218231 | X | | | | |
| ORL04 | 952 | 45,950 | SAMN41218232 | X | | | | |
| ORL05 | 953 | 155,426 | SAMN41218233 | X | X | X | X | X |
| ORL06 | 954 | 109,981 | SAMN41218234 | X | X | X | X | X |

| | | | | | | | | |
|---|---|---|---|---|---|---|---|---|
| ORL08 | 956 | 262,824 | SAMN41218236 | X | X | X | X | X |
| ORL09 | 957 | 180,747 | SAMN41218237 | X | X | X | X | X |
| ORL10 | 958 | 115,241 | SAMN41218238 | X | X | X | X | X |
| ORL12 | 960 | 53,472 | SAMN41218239 | X | | | | |
| ORL13 | 961 | 96,610 | SAMN41218240 | X | | | | |
| ORL14 | 962 | 104,556 | SAMN41218241 | X | X | X | X | X |
| ORL15 | 963 | 89,789 | SAMN41218242 | X | | | | |
| ORL16 | 964 | 55,596 | SAMN41218243 | X | | | | |
| ORL18 | 966 | 769,371 | SAMN41218245 | X | X | X | X | X |
| ORL20 | 968 | 594,887 | SAMN41218247 | X | X | X | X | X |
| ORL21 | 969 | 719,129 | SAMN41218248 | X | X | X | X | X |
| ORL22 | 970 | 146,658 | SAMN41218249 | X | X | X | X | X |
| ORL26 | 974 | 106,435 | SAMN41218250 | X | X | X | X | X |
| ORL32 | 980 | 9,410 | SAMN41218252 | X | | | | |
| ORL40 | 988 | 212,703 | SAMN41218255 | X | X | X | X | X |
| OTL01 | 1006 | 168,088 | SAMN41218256 | X | X | X | X | X |
| OTL03 | 1008 | 65,344 | SAMN41218257 | X | | | | |
| OTL04 | 1009 | 56,804 | SAMN41218258 | X | | | | |
| OTL05 | 1010 | 5,992 | SAMN41218259 | X | | | | |
| OTL06 | 1011 | 13,276 | SAMN41218260 | X | | | | |
| OTL07 | 1012 | 9,114 | SAMN41218261 | X | | | | |
| OTL08 | 1013 | 30,691 | SAMN41218262 | X | | | | |
| OTL09 | 1014 | 48,398 | SAMN41218263 | X | | | | |
| OTL15 | 1019 | 47,373 | SAMN41218265 | X | | | | |

| | | | | | | | | |
|---|---|---|---|---|---|---|---|---|
| OTL18 | 1022 | 219,867 | SAMN41218266 | X | X | X | X | X |
| OTL23 | 1027 | 58,007 | SAMN41218270 | X | | | | |
| OTL27 | 1031 | 76,544 | SAMN41218273 | X | | | | |
| OTL29 | 1033 | 634,961 | SAMN41218274 | X | X | X | X | X |
| OTL31 | 1035 | 416,726 | SAMN41218275 | X | X | X | X | X |
| OTL34 | 1038 | 28,824 | SAMN41218277 | X | | | | |
| RIL06 | 448 | 15,446 | SAMN41218183 | X | | | | |
| RIL18 | 460 | 557,749 | SAMN41218193 | X | X | X | X | X |
| RIL19 | 461 | 88,561 | SAMN41218194 | X | | | | |
| RIL21 | 463 | 25,829 | SAMN41218195 | X | | | | |
| RIL22 | 464 | 407,856 | SAMN41218196 | X | X | X | X | X |
| RIL23 | 465 | 43,165 | SAMN41218197 | X | | | | |
| RIL27 | 469 | 19,025 | SAMN41218199 | X | | | | |
| RIL29 | 471 | 314,614 | SAMN41218201 | X | X | X | X | X |
| RIL30 | 472 | 26,106 | SAMN41218202 | X | | | | |
| RIL31 | 473 | 51,275 | SAMN41218203 | X | | | | |
| RIL33 | 475 | 15,587 | SAMN41218204 | X | | | | |
| RTL07 | 489 | 133,294 | SAMN41218208 | X | X | X | X | X |
| RTL35 | 517 | 35,124 | SAMN41218227 | X | | | | |
| RTL36 | 518 | 278,193 | SAMN41218228 | X | X | X | X | X |

**Table S3:** Count and percentage of samples in the phylogenetic dataset and the population genomic data subset. RD = river delta, FR = fore reef.

| | | **Phylogenetic R30** | | | | | **Population Genomic R80A & R80S** | | | | | |
|---|---|---|---|---|---|---|---|---|---|---|---|---|
| **Clade** | **Environment** | **Fouha** | **Inarajan** | **Talofofo** | **Ritidian** | **Total** | **Fouha** | **Inarajan** | **Talofofo** | **Ritidian** | **Total** | |
| **I** | **RD** | 0 | 0 | 0 | 0 | **0** | 0 | 0 | 0 | 0 | **0** | |
| | **FR** | 1 | 1 | 0 | 1 | **3** | 1 | 1 | 0 | 0 | **2** | 67% |
| **II** | **RD** | 2 | 2 | 1 | 0 | **5** | 1 | 1 | 0 | 0 | **2** | 40% |
| | **FR** | 4 | 9 | 3 | 4 | **20** | 4 | 6 | 2 | 0 | **12** | 60% |
| **III** | **RD** | 0 | 0 | 0 | 0 | **0** | 0 | 0 | 0 | 0 | **0** | |
| | **FR** | 0 | 1 | 1 | 1 | **3** | 0 | 0 | 0 | 0 | **0** | |
| **IV** | **RD** | 0 | 0 | 0 | 0 | **0** | 0 | 0 | 0 | 0 | **0** | |
| | **FR** | 0 | 2 | 4 | 1 | **7** | 0 | 0 | 4 | 0 | **4** | 57% |
| **V** | **RD** | 27 | 13 | 18 | 0 | **58** | 18 | 7 | 15 | 0 | **40** | 69% |
| | **FR** | 2 | 3 | 0 | 1 | **6** | 1 | 1 | 0 | 1 | **3** | 50% |
| **VI** | **RD** | 0 | 0 | 0 | 0 | **0** | 0 | 0 | 0 | 0 | **0** | |
| | **FR** | 8 | 0 | 1 | 0 | **9** | 1 | 0 | 1 | 0 | **2** | 22% |
| **VII** | **RD** | 0 | 11 | 3 | 0 | **14** | 0 | 3 | 2 | 0 | **5** | 35% |
| | **FR** | 5 | 8 | 15 | 19 | **47** | 3 | 5 | 4 | 12 | **24** | 51% |
| **Total** | **RD** | 29 | 26 | 22 | 0 | **77** | 19 | 11 | 17 | 0 | **47** | 61% |
| | **FR** | 20 | 24 | 24 | 27 | **95** | 10 | 13 | 11 | 13 | **47** | 49% |

### Clade distribution among geographic populations

**<u>Table S4:</u>** Samples in each geographic location per clade.

| | I | II | III | IV | V | VI | VII | **SUM** |
|---|---|---|---|---|---|---|---|---|
| Fouha | 1 (2%) | 6 (12%) | 0 (0%) | 0 (0%) | 29 (59%) | 8 (16%) | 5 (10%) | **49 (28%)** |
| Talofofo | 0 (0%) | 4 (9%) | 1 (2%) | 4 (9%) | 18 (39%) | 1 (2%) | 18 (39%) | **46 (27%)** |
| Inarajan | 1 (2%) | 11 (22%) | 1 (2%) | 2 (4%) | 16 (32%) | 0 (0%) | 19 (38%) | **50 (29%)** |
| Ritidian | 1 (4%) | 4 (15%) | 1 (4%) | 1 (4%) | 1 (4%) | 0 (0%) | 19 (70%) | **27 (16%)** |

**<u>Table S5:</u>** Samples in each clade per geographic location.

| | I | II | III | IV | V | VI | VII |
|---|---|---|---|---|---|---|---|
| Fouha | 1 (33%) | 6 (24%) | 0 (0%) | 0 (0%) | 29 (45%) | 8 (89%) | 5 (8%) |
| Talofofo | 0 (0%) | 4 (16%) | 1 (33%) | 4 (57%) | 18 (28%) | 1 (11%) | 18 (30%) |
| Inarajan | 1 (33%) | 11 (44%) | 1 (33% | 2 (29%) | 16 (25%) | 0 (0%) | 19 (31%) |
| Ritidian | 1 (33%) | 4 (16%) | 1 (33%) | 1 (14%) | 1 (2%) | 0 (0%) | 19 (31%) |
| SUM | **3 (2%)** | **25 (15%)** | **3 (2%)** | **7 (4%)** | **64 (37%)** | **9 (5%)** | **61 (35%)** |

### Color phenotypes within and among clades

**Table S6:** Percentage of samples per color morphotype belonging to each clade.

| | I | II | III | IV | V | VI | VII | **SUM** |
|---|---|---|---|---|---|---|---|---|
| Blue | 0 (0%) | 0 (0%) | 0 (0%) | 1 (33%) | 0 (0%) | 0 (0%) | **2 (67%)** | **3** |
| Purple | 2 (13%) | 3 (19%) | 0 (0%) | 0 (0%) | 0 (0%) | 5 (31%) | **6 (38%)** | **16** |
| Green | 0 (0%) | 0 (0%) | 0 (0%) | 0 (0%) | 7 (37%) | 0 (0%) | **12 (63%)** | **19** |
| Yellow | 1 (2%) | 11 (18%) | 1 (2%) | 2 (3%) | 21 (35%) | 0 (0%) | **24 (40%)** | **60** |
| Cream | 0 (0%) | 0 (0%) | 0 (0%) | 0 (0%) | **2 (50%)** | 0 (0%) | **2 (50%)** | **4** |
| Brown | 0 (0%) | 8 (13%) | 2 (3%) | 3 (5%) | **32 (51%)** | 5 (8%) | 13 (21%) | **63** |

**Table S7:** Percentage of samples in each clade with each color morphotype.

| | I | II | III | IV | V | VI | VII | **SUM** | **Clades** |
|---|---|---|---|---|---|---|---|---|---|
| Blue | 0 (0%) | 0 (0%) | 0 (0%) | 1 (17%) | 0 (0%) | 0 (0%) | 2 (3%) | **3** | **2** |
| Purple | **2 (67%)** | 3 (14%) | 0 (0%) | 0 (0%) | 0 (0%) | **5 (50%)** | 6 (10%) | 16 | 4 |
| Green | 0 (0%) | 0 (0%) | 0 (0%) | 0 (0%) | 7 (11%) | 0 (0%) | 12 (20%) | 19 | 2 |
| Yellow | 1 (33%) | **11 (50%)** | 1 (33%) | 2 (33%) | 21 (34%) | 0 (0%) | **24 (41%)** | 60 | 6 |
| Cream | 0 (0%) | 0 (0%) | 0 (0%) | 0 (0%) | 2 (3%) | 0 (0%) | 2 (3%) | 4 | 2 |
| Brown | 0 (0%) | 8 (36%) | **2 (67%)** | **3 (50%)** | **32 (52%)** | **5 (50%)** | 13 (22%) | 63 | 6 |
| **SUM** | **3** | **22** | **3** | **6** | **62** | **10** | **59** | **165** | |
| **Colors** | **2** | **3** | **2** | **3** | **4** | **2** | **6** | | |

### AMOVAs

**Table S8:** AMOVA analyses of the three largest clades (II, V, VII).
SD = standard deviation; CI = confidence intervals.
Significance was tested using 999 permutations. Standard deviations of F-statistics were obtained through jackknifing over loci. 95% confidence intervals of F-statistics were obtained through bootstrapping over loci.

A) AMOVA of all 7 clades (96 samples). Both clones were removed, but OIL35 from clade III was included.

| Source of Variation | % Variation | F-Stat | F-value | SD | CI (2.5%) | CI (97.5%) | P-value | F'-value |
|---|---|---|---|---|---|---|---|---|
| **Within Individual** | 0.324 | F_it | 0.676 | 0.007 | 0.663 | 0.689 | -- | -- |
| **Within Population** | 0.065 | F_is | 0.167 | 0.006 | 0.155 | 0.179 | **0.001** | -- |
| **Among Populations** | **0.611** | F_st | 0.611 | 0.008 | 0.596 | 0.626 | **0.001** | 0.639 |

B) AMOVA of the three largest clades (II, V, VII); 86 samples.

| Source of Variation | % Variation | F-Stat | F-value | SD | CI (2.5%) | CI (97.5%) | P-value | F'-value |
|---|---|---|---|---|---|---|---|---|
| **Within Individual** | 0.316 | F_it | 0.684 | 0.008 | 0.669 | 0.698 | -- | -- |
| **Within Population** | 0.059 | F_is | 0.158 | 0.007 | 0.145 | 0.171 | **0.001** | -- |
| **Among Populations** | **0.624** | F_st | 0.624 | 0.008 | 0.608 | 0.640 | **0.001** | 0.652 |

C) AMOVA of the three largest clades (II, V, VII) with all clades subsampled to 14 samples to account for uneven sample sizes.

| Source of Variation | % Variation | F-Stat | F-value | SD | CI (2.5%) | CI (97.5%) | P-value | F'-value |
|---|---|---|---|---|---|---|---|---|
| **Within Individual** | 0.268 | F_it | 0.732 | 0.007 | 0.718 | 0.746 | -- | -- |
| **Within Population** | 0.052 | F_is | 0.163 | 0.009 | 0.146 | 0.180 | **0.001** | -- |
| **Among Populations** | **0.680** | F_st | 0.680 | 0.008 | 0.665 | 0.695 | **0.001** | 0.712 |

**<u>Table S9:</u>** Clade V geography AMOVA excluded the sole Ritidian sample (ORL07), and included both river delta and fore reef samples at each geographic location.
Fouha: n=19
Inarajan: n=8
Talofofo: n=15

| Source of Variation | % Variation | F-Stat | F-value | SD | CI (2.5%) | CI (97.5%) | P-value | F'-value |
|---|---|---|---|---|---|---|---|---|
| **Within Individuals** | 0.829 | F_it | 0.171 | 0.002 | 0.167 | 0.175 | -- | -- |
| **Within Populations** | 0.167 | F_is | 0.167 | 0.002 | 0.163 | 0.171 | **0.001** | -- |
| **Among Populations** | **0.005** | F_st | 0.005 | 0.000 | 0.004 | 0.005 | **0.001** | 0.005 |

SD = standard deviation; CI = confidence intervals

**<u>Table S10:</u>** Clade VII geography AMOVA excluded the Fouha population (n=3) due to low sample size, and included both river delta and fore reef samples at each geographic location.Clade VII geography AMOVA.
Inarajan: n=8
Ritidian: n=12
Talofofo: n=6

| Source of Variation | % Variation | F-Stat | F-value | SD | CI (2.5%) | CI (97.5%) | P-value | F'-value |
|---|---|---|---|---|---|---|---|---|
| **Within Individuals** | 0.853 | F_it | 0.147 | 0.003 | 0.142 | 0.152 | -- | -- |
| **Within Populations** | **0.143** | F_is | 0.143 | 0.003 | 0.138 | 0.149 | **0.001** | -- |
| **Among Populations** | **0.004** | F_st | 0.004 | 0.001 | 0.003 | 0.006 | **0.001** | 0.005 |

SD = standard deviation; CI = confidence intervals

### Pairwise $F_{ST}$

Top diagonal: p-value
Bottom diagonal: pairwise $F_{ST}$

**<u>Table S11:</u>** Clade V pairwise $F_{ST}$ excluded the sole Ritidian sample (ORL07), and included both river delta and fore reef samples at each geographic location.
Fouha: n=19
Inarajan: n=8
Talofofo: n=15

| | Fouha | Inarajan | Talofofo |
|---|---|---|---|
| Fouha | -- | 0.521 | **0.001** |
| Inarajan | 0.000 | -- | 0.088 |
| Talofofo | 0.008 | 0.003 | -- |

**<u>Table S12:</u>** Clade VII pairwise $F_{ST}$ excluded the Fouha population (n=3) due to low sample size, and included both river delta and fore reef samples at each geographic location.
Inarajan: n=8
Ritidian: n=12
Talofofo: n=6

| | Inarajan | Ritidian | Talofofo |
|---|---|---|---|
| Inarajan | -- | 0.068 | 0.316 |
| Ritidian | 0.005 | -- | 0.149 |
| Talofofo | 0.001 | 0.005 | -- |

**Table S13:** Symbiont Read counts across samples containing symbiont reads.

| Sample | *Symbiodinium* | *Breviolum* | *Cladocopium* | *Durusdinium* |
|---|---|---|---|---|
| OFL07 | 0 | 0 | 16 | 0 |
| OFL30 | 0 | 0 | 3 | 0 |
| OFL32 | 0 | 0 | 1 | 0 |
| OFL33 | 0 | 0 | 2 | 0 |
| OFL36 | 0 | 0 | 18 | 0 |
| OFL04 | 0 | 0 | 56 | 0 |
| OIL03 | 0 | 0 | 1 | 0 |
| OIL05 | 0 | 0 | 3 | 0 |
| OIL11 | 0 | 0 | 7 | 0 |
| OIL19 | 0 | 0 | 139 | 0 |
| OIL21 | 0 | 0 | 29 | 0 |
| OIL26 | 0 | 0 | 22 | 0 |
| OIL32 | 0 | 0 | 34 | 0 |
| OIL33 | 0 | 0 | 4 | 0 |
| ORL05 | 0 | 0 | 2 | 0 |
| ORL08 | 0 | 0 | 4 | 0 |
| ORL09 | 0 | 0 | 1 | 0 |
| ORL20 | 0 | 0 | 3 | 0 |
| OTL01 | 0 | 0 | 3 | 0 |
| OTL10 | 0 | 0 | 5 | 0 |
| OTL20 | 0 | 0 | 6 | 0 |
| OTL29 | 0 | 0 | 5 | 0 |

| | | | | |
|---|---|---|---|---|
| OTL31 | 0 | 0 | 4 | 1 |
| OTL36 | 0 | 0 | 3 | 0 |
| RFL08 | 0 | 0 | 9 | 0 |
| RFL14 | 0 | 0 | 4 | 0 |
| RFL30 | 0 | 0 | 3 | 0 |
| RFL31 | 0 | 0 | 10 | 0 |
| RFL32 | 0 | 0 | 2 | 0 |
| RFL33 | 0 | 0 | 7 | 0 |
| RFL36 | 0 | 0 | 10 | 0 |
| RIL07 | 0 | 0 | 3 | 0 |
| RIL09 | 0 | 1 | 1 | 0 |
| RIL11 | 0 | 0 | 62 | 0 |
| RIL17 | 0 | 0 | 2 | 0 |
| RIL18 | 0 | 0 | 9 | 0 |
| RIL29 | 0 | 0 | 3 | 1 |
| RTL01 | 0 | 1 | 1 | 0 |
| RTL09 | 0 | 0 | 1 | 0 |
| RTL11 | 0 | 0 | 9 | 0 |
| RTL15 | 0 | 0 | 11 | 0 |
| RTL18 | 0 | 0 | 9 | 0 |
| RTL26 | 0 | 0 | 11 | 0 |
| RTL36 | 0 | 0 | 7 | 0 |

### Phylogenomics

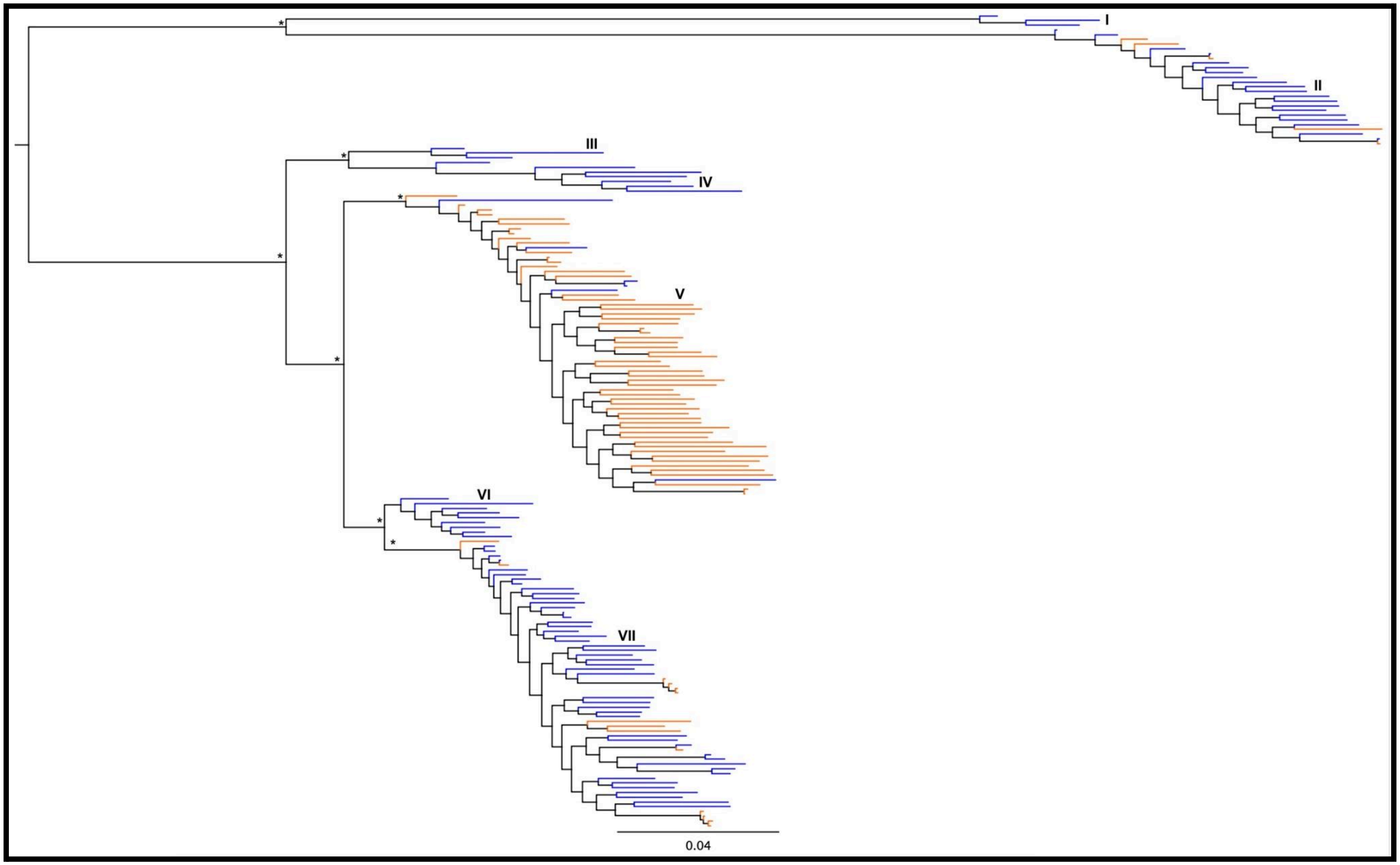

**Figure S1:** IQTree phylogenetic tree using 22,982 loci and 110,252 concatenated SNPs (lnL= -1,688,552). Node support is based on 1000 bootstraps replicates. Tip colors indicate the environmental origin of each sample. Asterisks indicate 100% bootstrap support.

### Clonality

**Figure S2:** Clonality IBS dendrograms for clade II (top left), IV (top right), V (bottom left), and VII (bottom right). Technical replicates are indicated in circles, while clones are indicated by red asterisks.

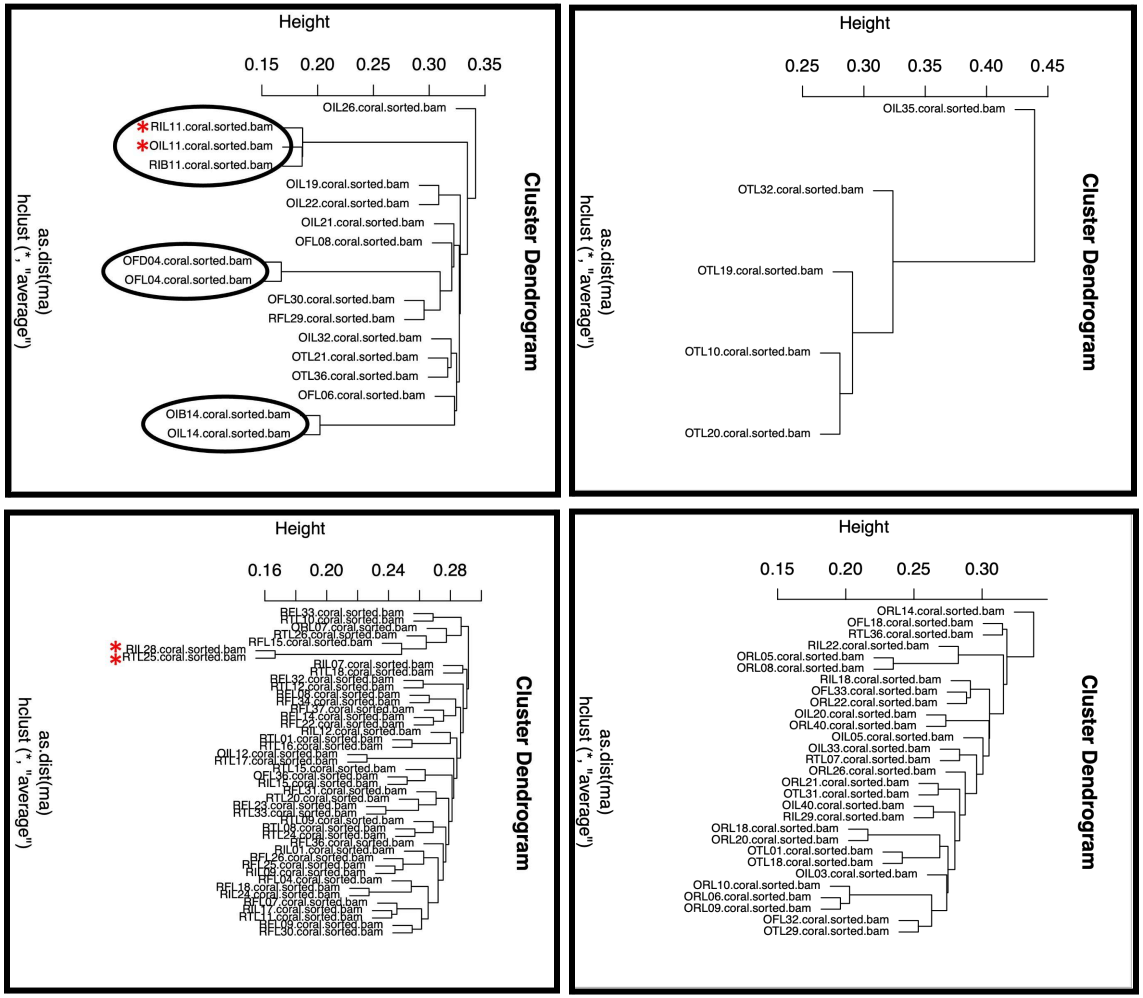

### NGSAdmix plots

**Figure S3:** K=2-8 genomic dataset NGSAdmix plots. All y-axes refer to admixture proportions.

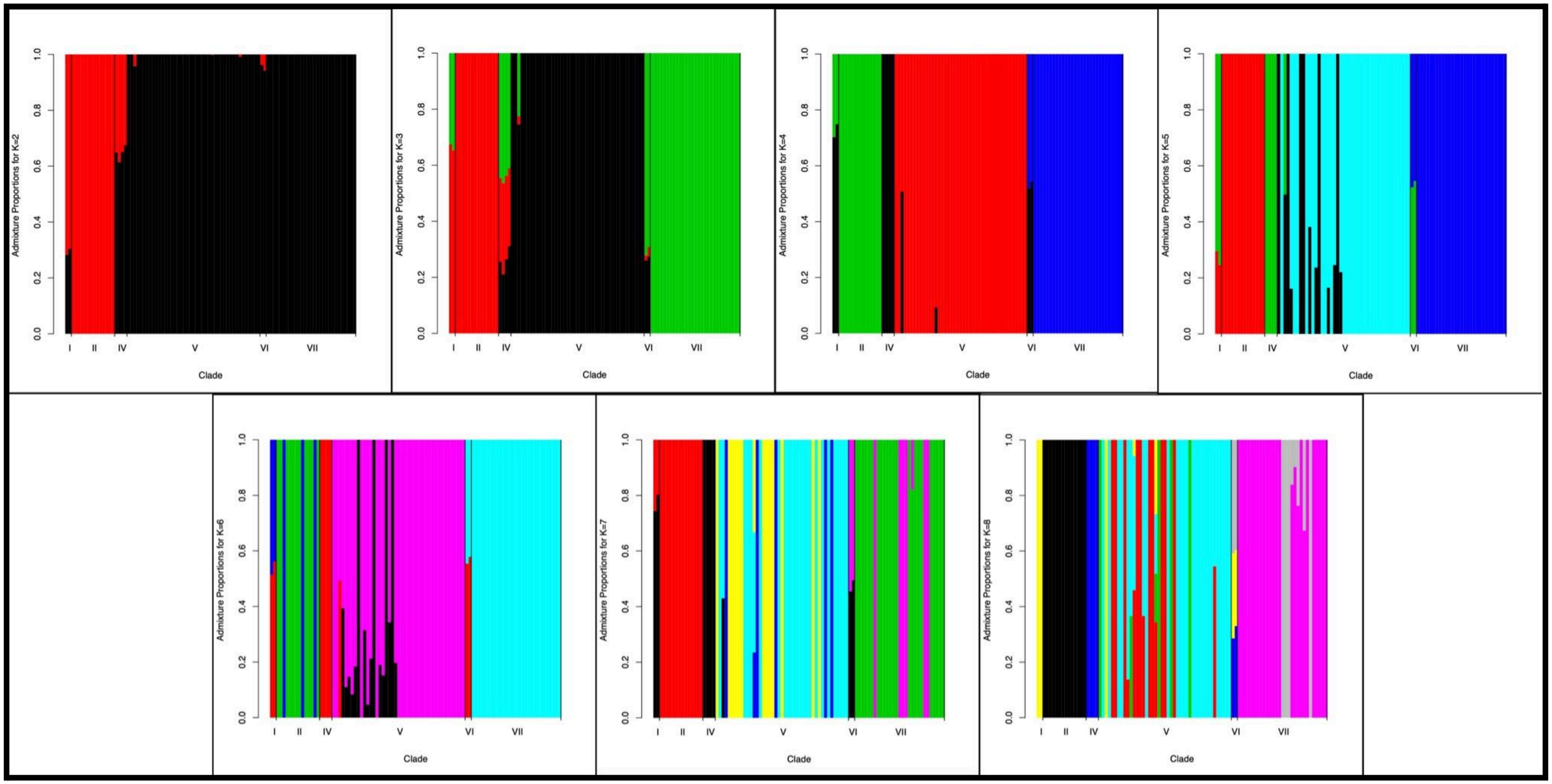

**Figure S4:** K=2-5 admixture plot for Clades I-II-IV. All y-axes refer to admixture proportions. Most importantly, none of these analyses supports a mixed ancestry of Clade I samples. While analyses with K2 and K3 do not distinguish Clades I and IV, analyses with K = 4 and 5, clearly distinguish them as separate clusters.

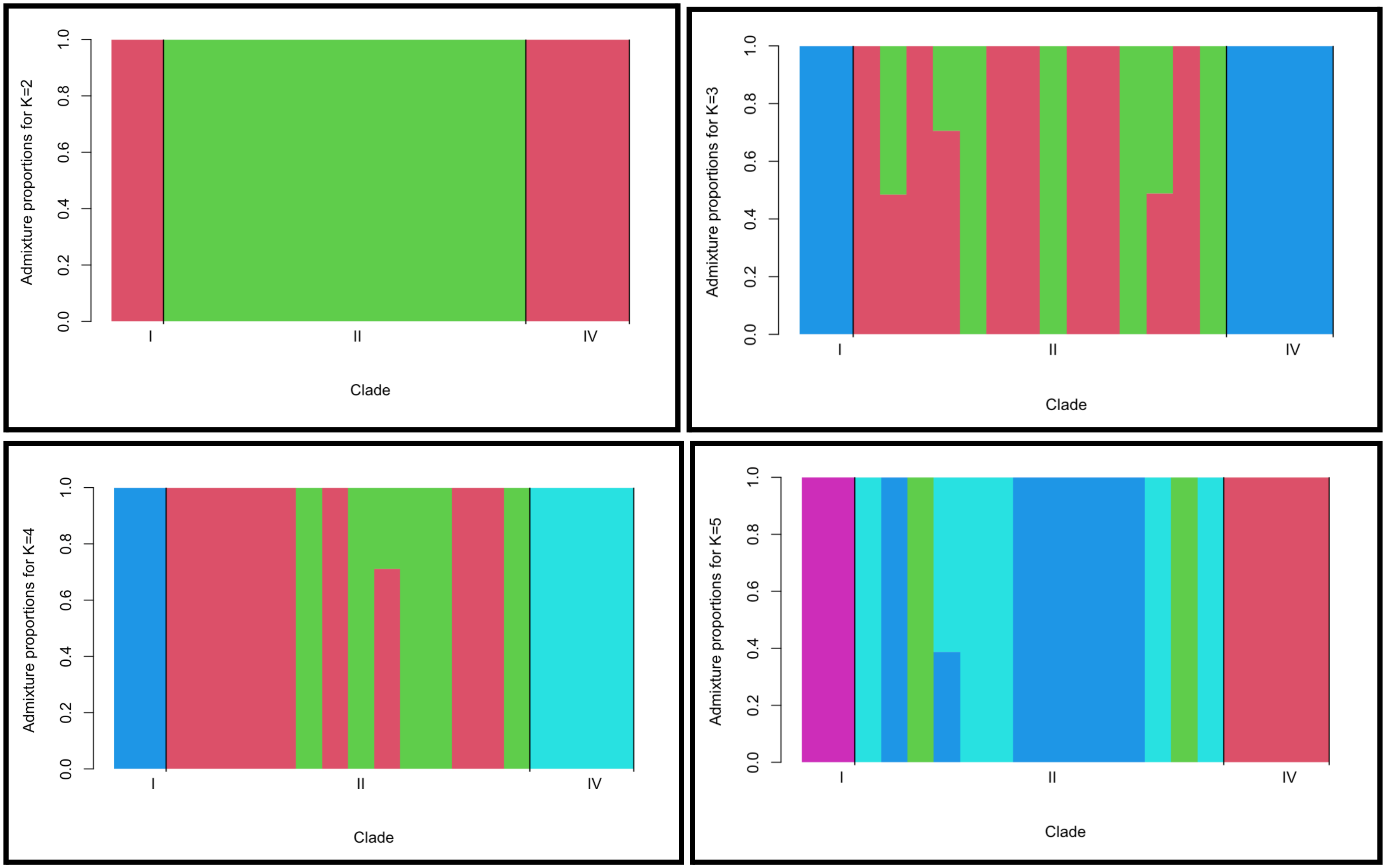

**Figure S5:** K=2-5 admixture plot for Clades IV - VII. All y-axes refer to admixture proportions.
All four analyses indicate that both Clade VI specimens in this dataset have a mixed ancestry. Analyses with $K \geq 3$ indicate that they may be hybrids between clades IV and VII (see also Fig. S5).

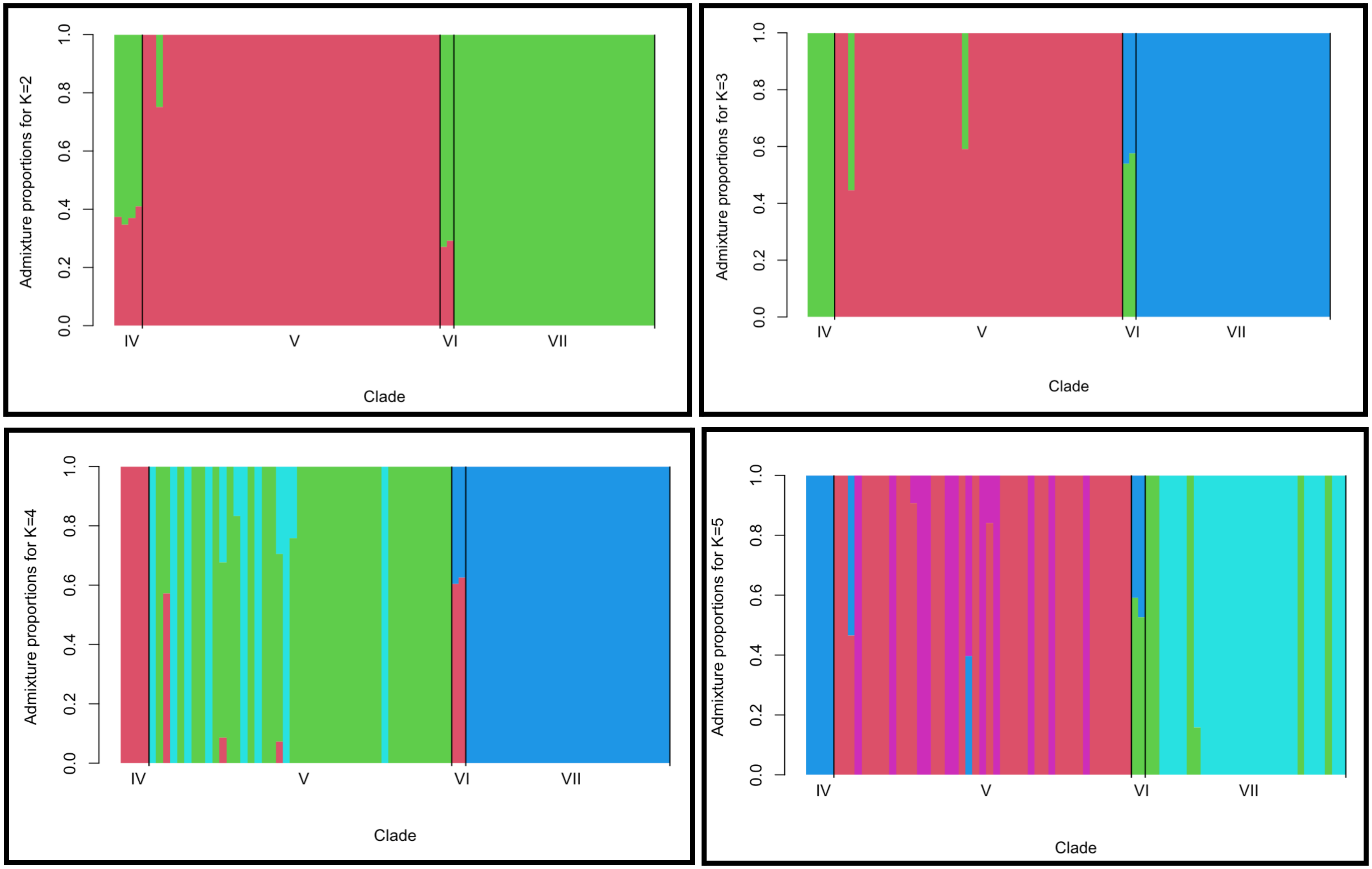

**Figure S6:** K=2-5 admixture plot for Clades IV, VI, & VII. All y-axes refer to admixture proportions. All four analyses indicate that both clade VI specimens in this dataset have a mixed ancestry and may be hybrids between clades IV and VII .

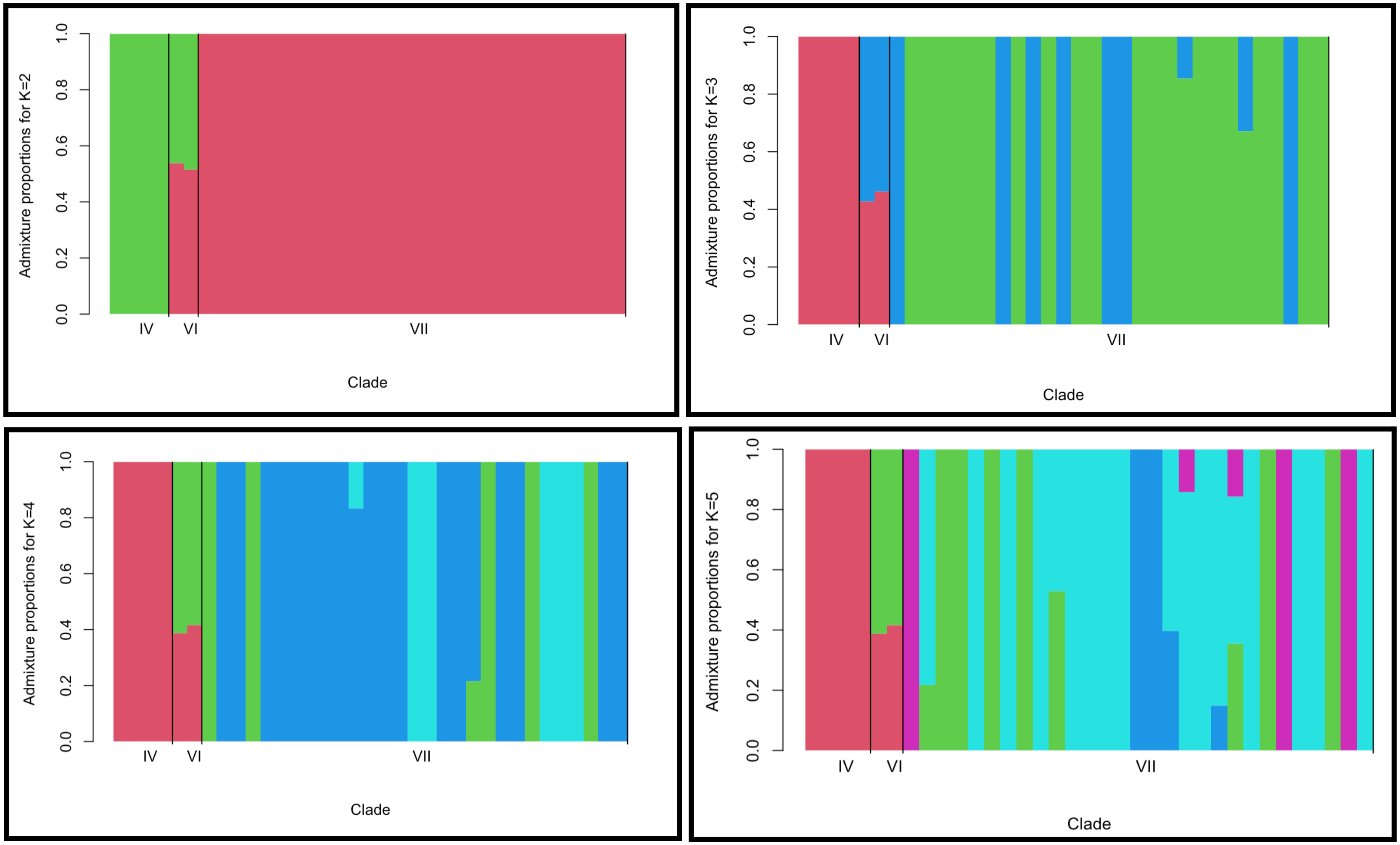

**Figure S7:** Intra-specific admixture plots for Clade V (top) and Clade VII (bottom) for K=2-4. All y-axes refer to admixture proportion. Results indicate no clear population genetic structure among populations.

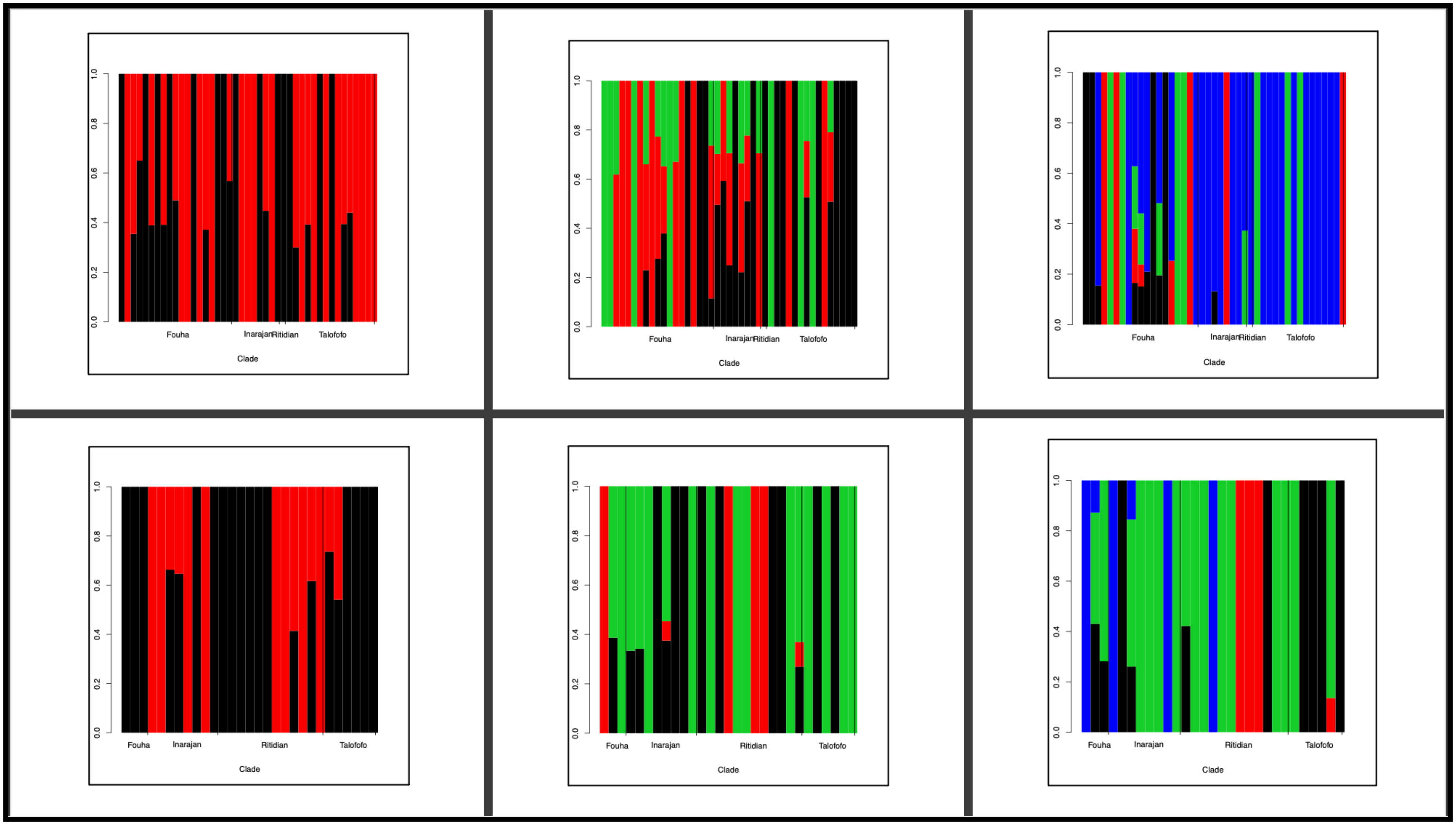
